# The obesity-linked peptide SP16 regulates adipocytes through GIP and insulin receptor

**DOI:** 10.1101/2025.06.17.660082

**Authors:** Thi My Hanh Ngo, Rakesh Santhanam, Daniel P. Teufel, Alec Dick, Daniel Lam, Holger Klein, Andreas-David Brunner, Mafalda M.A. Pereira, Alexander Bartelt, Anton Pekcec, Maude Giroud

## Abstract

Obesity has emerged as a global epidemic and represents a major public health concern due to its profound implications for metabolic diseases. While recent pharmacological interventions primarily aim to reduce food intake, enhancing energy expenditure in adipose tissue offers a promising complementary approach. In this study, we identify that SP16, a synthetic α1- antitrypsin derived peptide, is a novel GIP (gastric inhibitory peptide) receptor agonist. SP16 promotes lipolysis via the cAMP pathway, resulting in increased mitochondrial oxygen consumption in adipocytes. Furthermore, SP16 binds to the insulin receptor, and at high concentrations, it attenuates insulin receptor signaling. In diet-induced obese (DIO) mice, acute SP16 treatment improved glucose clearance, elevated circulating free fatty acids, and decreased leptin levels. *Ex vivo* analyses of epididymal white adipose tissue corroborated these findings, demonstrating enhanced lipolysis in SP16-treated mice. In conclusion, SP16 emerges as a novel lipolytic peptide with beneficial metabolic effects.

**Graphical abstract:** 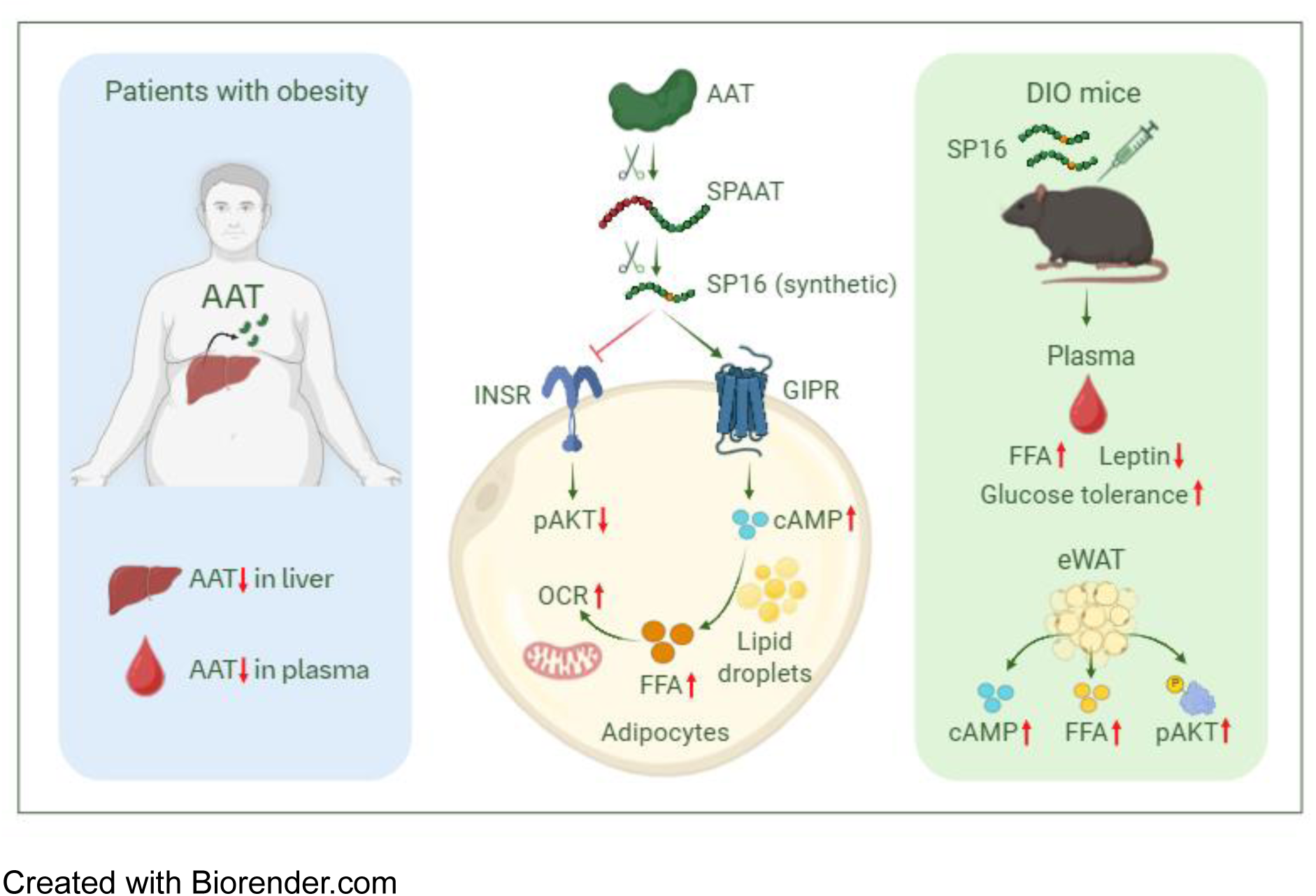

Created with Biorender.com

## Introduction

Obesity is a multifaceted and chronic disease characterized by a dysregulation of energy homeostasis leading to excessive accumulation of adipose tissue^1^. It is associated with various health complications, including type 2 diabetes, cardiovascular diseases, cancer, and musculoskeletal disorders^1^. The detrimental impact of obesity extends beyond physical health, contributing to psychological distress and economic burden, highlighting the urgent need for effective therapeutic intervention^2^. Therapeutics that influence energy intake (EI), energy expenditure (EE), or both are necessary to support patients living with obesity^3^.

Peptide hormones are crucial regulators of systemic energy balance. The identification of glucagon-like peptide 1 (GLP-1), an incretin hormone, revolutionized the development of the first generation of anti-obesity therapies by its pivotal role in regulating food intake and glucose metabolism^4^. New generations of anti-obesity drugs such as dual agonists include a GIPR agonist component. GIP has demonstrated the ability to stimulate glucose-dependent insulin secretion, induce weight loss, and improve glucose tolerance^5–7^.

Preclinical studies shed light on mechanisms that enhance adipocyte metabolism, leading to increased thermogenesis and EE, ultimately promoting weight loss. Therefore, targeting adipocytes holds promise for advancing current obesity treatments. Several peptides, including GIP, GLP-1, brown adipose tissue (BAT)-secreted peptide 1 along with mitochondria-derived peptides (humanin, small humanin-like peptide 2, MOTS-c) have been identified and characterized for their ability to promote adipocyte thermogenesis, enhance lipid oxidation, and prevent weight gain induced by a high-fat diet (HFD)^8–10^. Furthermore, it has been demonstrated that activating GIPR in adipocytes leads to increased fat thermogenesis and EE via promoting energy wasting through calcium futile cycle, thereby promoting weight loss in obese mice^11^.

Proteins implicated in obesity have the potential to modulate adipocyte metabolism. One such protein, α1-antitrypsin (AAT), primarily produced in the liver and to a lesser extent expressed in macrophages, pancreatic islets, and endothelial cells, is a broad-spectrum protease inhibitor^12^. Its concentration can increase from 1-2 g/L up to four-fold during the acute phase of inflammation^12^. Its primary function is to inhibit neutrophil elastase, thereby protecting tissues, particularly the lungs from proteolytic damage^12^. AAT deficiency is associated with an elevated risk of chronic obstructive pulmonary disease and liver disease and can be managed through augmentation therapy with purified AAT^12^. Additionally, emerging evidence also implicates AAT in metabolic disorders, including obesity. Studies in mice have demonstrated that both liver and plasma levels of AAT are significantly downregulated in ob/ob mice compared to wild-type mice^13,14^. Additionally, overexpression of human AAT in mice has been shown to protect against HFD-induced obesity, insulin resistance, fatty liver, and inflammation^14^. AAT can be cleaved at carboxyl-terminal end by proteases, such as metalloproteinases, into peptides known as carboxyl-terminal peptides of AAT (CAAPs)^15^. Several CAAPs, including C21, C36, C42, and C44 (also known as SPAAT - Short Peptide AAT), have been detected in various human tissues and fluids and identified as biomarkers for inflammation and preeclampsia activity^15^. Multiple AAT-derived peptides have been developed to explore their therapeutic potential^16–18^. Among these, SP16 is a synthetic 17- amino acid peptide corresponding to residues 388-404 of AAT, in which methionine is substituted with norleucine to enhance proteolytic stability.

While the association between AAT and obesity is well described, peptides derived from AAT have not been investigated in *in vitro* and *in vivo* models of obesity. Therefore, in this study, we examined the beneficial effects of an AAT-derived peptide on lipid metabolism in adipocytes and glucose metabolism in diet-induced obesity (DIO) mice. We also investigated the signaling pathway as well as receptor regulating its bioactivity in adipocytes.

## Results

### SP16 is a peptide associated with obesity

Genetic associations with obesity and related endophenotypes suggest involvement in key regulatory mechanisms. In this study, we specifically investigate the role of peptides derived from these associated proteins in modulating adipocyte metabolism. To identify such candidates, we selected the top 20 genes associated with body weight based on the HuGE (Human Genetic Evidence) score from the Common Metabolic Diseases Knowledge Portal (https://hugeamp.org) (Fig. 1a). Subsequently, we examined whether the encoded proteins of these genes had annotated peptides in the Uniprot database. Our analysis revealed that only two proteins, ProSAAS (coded by *PCSK1* gene) and AAT (encoded by *SERPINA1* gene), are predicted to be cleaved into peptides, with ProSAAS yielding seven peptides and AAT producing one peptide, namely SPAAT (Supplementary Table 1). SPAAT is a 44-amino acid peptide derived from the C-terminal end of AAT (residues 388 to 418). Moreover, *Pcsk1* and *Serpina1* were identified as orthologous protein-coding genes retrieved from a mouse peptidomic study conducted by Madsen *et al*.^19^. In this analysis, the authors identified 157,857 peptides derived from 1,229 proteins across seven tissues from models of obese and diabetic mice^19^. We further compared the eight peptides retrieved from ProSAAS and AAT to the healthy human peptidomic dataset obtained from Arapidi *et al.*^20^. SPAAT and one peptide derived from ProSAAS but not annotated in Uniprot were detected in the human plasma peptidome. To conclude, among the eight Uniprot-annotated peptides, only SPAAT was detected (Fig. 1a). To investigate the association between AAT and obesity, we conducted proteomic analyses on liver samples from lean and DIO mice. The *Serpina1* gene family in mice contains six members, *Serpina1a* to *Serpin1f*. According to Uniprot.org, five *Serpina1* genes (from *Serpina1*a to *Serpina1*e) are expressed in liver, whereas *Serpina1f* is predominantly expressed in epididymis. Our proteomic results indicated that only SERPINA1E is significantly downregulated in both liver and plasma of obese mice compared to lean mice with a strong co-regulation across plasma and liver tissue (R = 0.88) (Fig. 1b, 1c, 1d). The proteomic result was confirmed at RNA level via qPCR with *Serpina1e* expression significantly downregulated in the liver of DIO mice (Fig. 1b). On the other hand, no significant differences in the mRNA levels of *Serpina1a, Serpina1b, Serpina1c*, and *Serpina1d* between lean and DIO mice were observed (Supplementary Fig. 1). Additionally, the hepatic marker of lipid accumulation *Cidea*, was considerably upregulated in response to HFD feeding (Fig. 1b). In humans, SERPINA1 encodes a single AAT isoform. Notably, AAT expression and function are reduced in obese individuals, and AAT is being explored as a potential therapeutic target for metabolic disorders^14,21^.

**Fig. 1:**
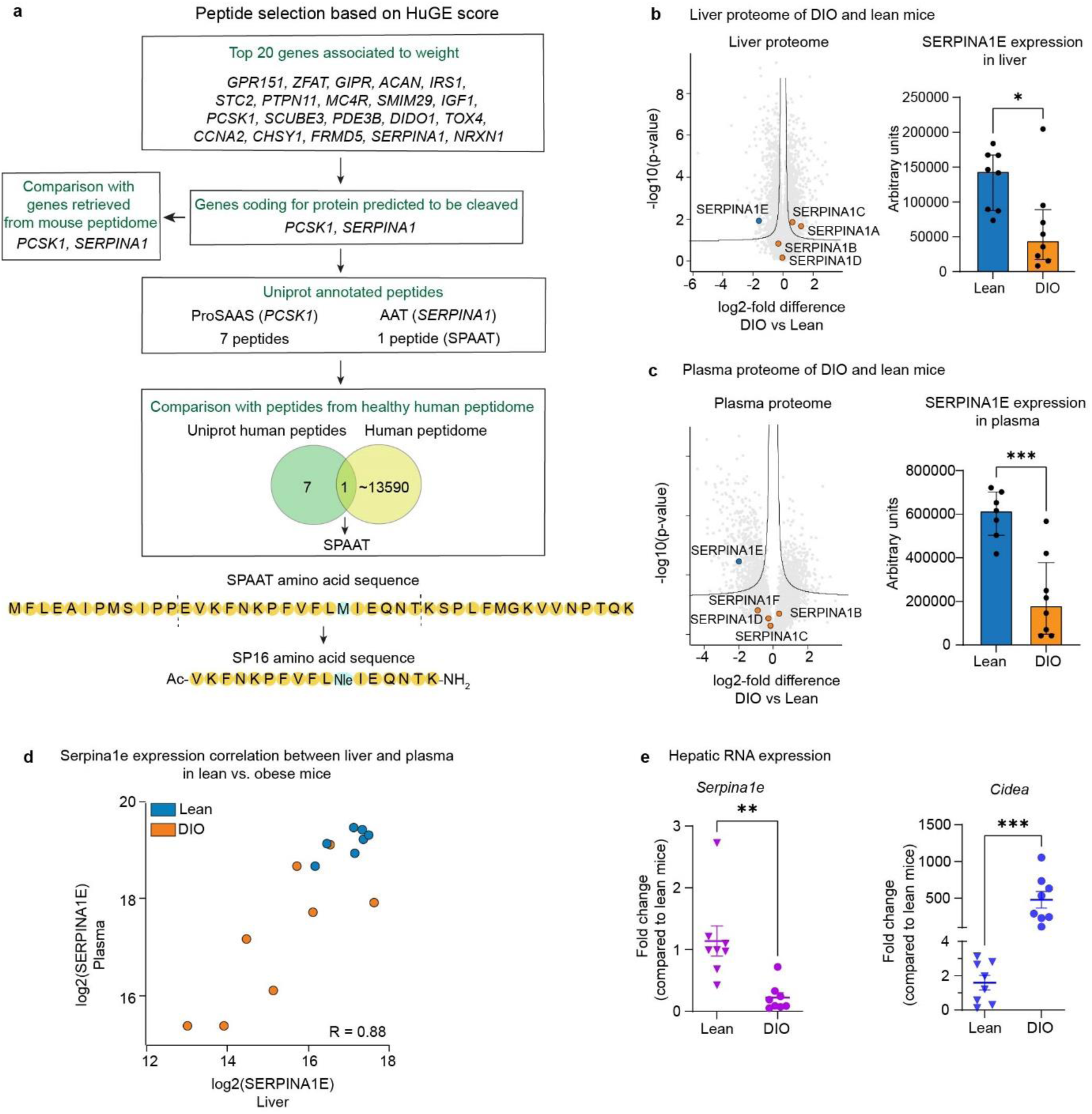
**Identification of AAT derived peptides with genetic association to obesity**. Identification cascade for peptides with genetic association to obesity (a), differential expression analysis of SERPINA1 family between lean and DIO mice in the liver (b), differential expression analysis of SERPINA1 family between lean and DIO mice in the plasma (c) correlation of differential expression analysis of Serpina1e between liver and plasma in lean and DIO mice (d), mRNA expression of *Serpina1e* and *Cidea* gene in liver of lean and DIO mice (b), HuGE: human genetic evidence. *PCSK1*: proprotein convertase 1, AAT: α-1 antitrypsin, SPAAT: short peptide AAT.

AAT is a serpin protein known to undergo cleavage into peptides at its C-terminal end^15^. While some of these peptides, including SPAAT, have been identified as biomarkers for sepsis, their precise roles and mechanisms of action remain largely unknown^15^. SP16, a synthetic peptide derived from SPAAT, has demonstrated its bioactivity in preventing LPS-induced inflammation and was already tested in phase 1 clinical trial for acute myocardial infarction and neuropathic pain^18,22^ (patent number WO2014/197524, US2019/0329952). SP16 was described in the literature as an agonist of LRP1^18^. Since this receptor is ubiquitously expressed, including in the adipose tissue^23^, we further investigated whether AAT, SPAAT or SP16 exhibits its bioactivity in an adipocyte model, shedding light on its potential implications in the context of adipose tissue.

### SP16 induces cAMP-dependent lipolysis in human adipocytes

AAT is described to regulate adipocytes metabolism^24^. Therefore, we tested the ability of AAT and AAT-derived peptides (SPAAT and SP16) to promote lipolysis on model of human white adipocytes, using human multipotent adipose-derived stem cells (hMADS). Lipolysis assays were conducted, with isoprenaline used as a positive control (differentiation method depicted in Fig. 2a). SP16 is able to induce FFA release in white hMADS cells (Fig. 2b), whereas ATT and SPAAT don’t (Supplementary Fig. 2a). To determine whether SP16 also exerts bioactivity in other adipocyte models, we extended our analysis to brite hMADS (Fig. 2e-2f) cells and differentiated human stromal vascular fraction (hSVF) cells (Fig. 2g-2h). Our findings revealed that SP16, but not SP16 negative control (SP16 NC), effectively induces lipolysis, with EC50 values of 7.43 µM, 18 µM, and 6.5 µM observed for white hMADS, brite hMADS, and white hSVF, respectively (Fig. 2a, 2f, 2h). To elucidate the signaling pathway underlying the lipolytic effect of SP16, we utilized an adenylyl cyclase inhibitor (SQ22536) and measured FFA release. Isoprenaline and BNP (brain natriuretic peptide) were used as controls, as isoprenaline induces lipolysis via cAMP while BNP acts through cGMP. Release of FFAs induced by both isoprenaline and SP16, is inhibited by SQ22536, while the release of FFAs induced by BNP remains unaffected (Fig. 2c). This finding is further confirmed by direct ELISA measurement of cellular cAMP levels (Fig. 2d). In conclusion, the lipolytic effect of SP16 is mediated through the cAMP pathway.

**Fig. 2:**
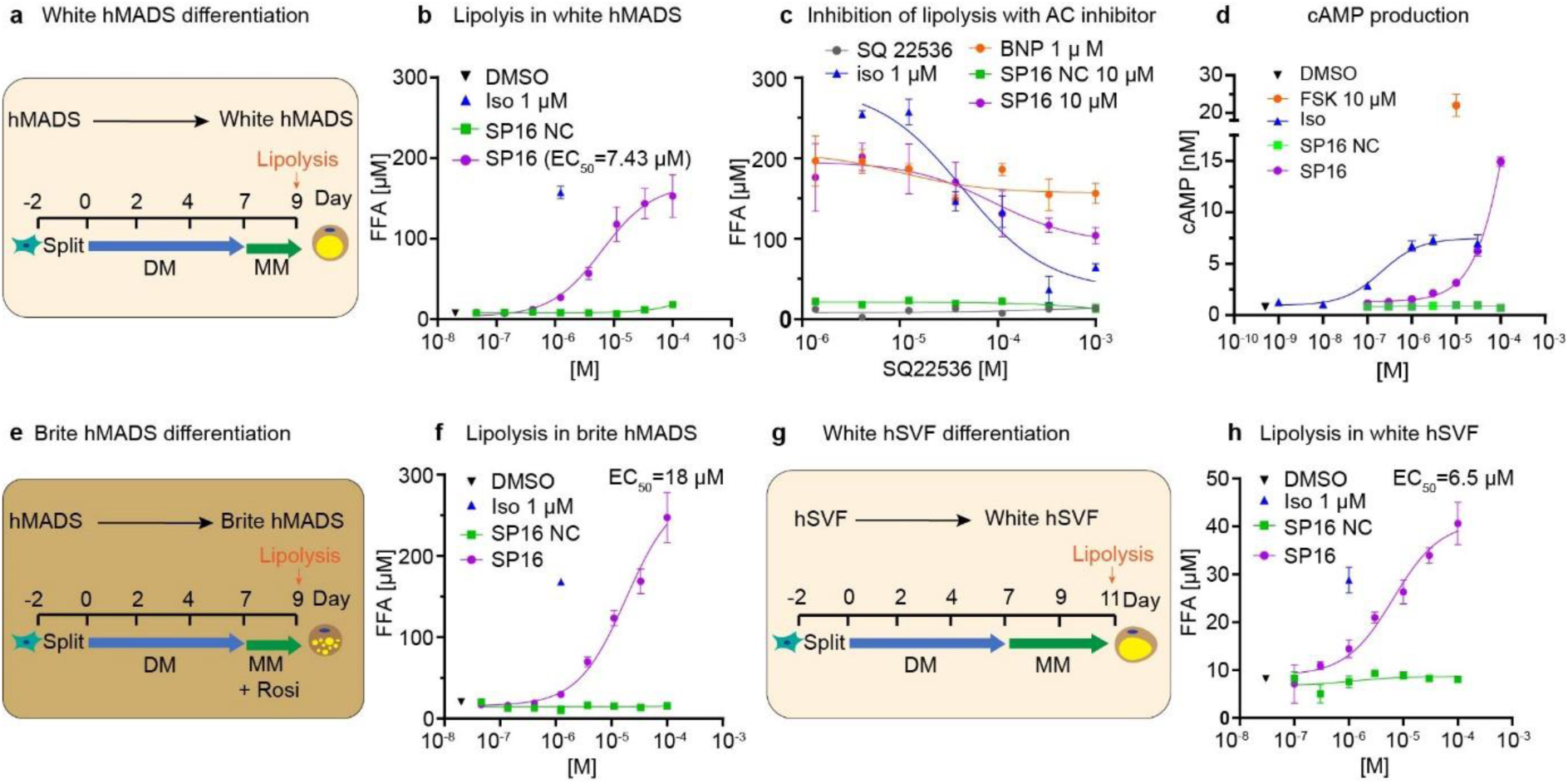
SP16 induces cAMP-dependent lipolysis in human adipocytes. Differentiation schemes and protocol for white hMADS (a), brite hMADS (e), and white hSVF (g), SP16-driven FFA release in white hMADS (b), brite hMADS (f), differentiated hSVF (h), inhibition of FFA release in white hMADS pretreated with SQ22636 (adenylyl cyclase inhibitor) (c), SP16-induced cAMP concentration in white hMADS (d). Data are mean ± SEM, n=4-8. hMADS: human multipotent adipose derived stem cells, DM: differentiation medium, MM: maintenance medium, Rosi: rosiglitazone, hSVF: human stromal vascular fraction, iso: isoprenaline, AC: adenylyl cyclase, FSK: forskolin, BNP: B-type natriuretic peptide, SP16 NC: SP16 negative control.

### SP16 promotes mitochondrial oxygen consumption through lipid oxidation

To evaluate whether SP16-induced lipolysis is linked to a higher oxidation of FFAs, seahorse assays were conducted (Fig. 3a). Acute treatment of 10 µM SP16 in white hMADS demonstrated a higher maximal respiration, ATP production, coupling efficiency, and spare respiratory capacity compared to the SP16 NC (Fig. 3b, 3c). Proton leak does not significantly differ between the SP16 and SP16 NC groups, indicating that the increased basal and maximal respiration observed was primarily attributed to β-oxidation of fatty acids as substrates. This finding was in line with an unchanged expression of UCP1 mRNA levels after SP16 treatment (Fig. 3f). Furthermore, both 24-h and 3 x 24-h treatment of SP16 does not result in an increase in OCR, suggesting that the effect of SP16 on mitochondrial respiration is rather acute (Fig. 3a, Supplementary Fig. 2b, 2c). In the fatty acid oxidation test, SP16 exhibits higher maximal respiration and ATP production in the presence of bovine serum albumin (BSA) compared to the SP16 NC (Fig. 3d). However, when palmitate-BSA was introduced, OCR increased but no significant difference was observed between the SP16- treated group and the SP16 NC-treated group (Fig. 3d, 3e). This finding suggests that SP16 enhances mitochondrial oxygen consumption in a similar manner to palmitate-BSA but does not have an additive effect. Forskolin treatment, used as a positive control, demonstrates an additional improvement in maximal respiration along with palmitate-BSA (Supplementary Fig. 2d), which is not observed in hMADS cells treated with SP16. The mitochondrial respiration is impaired by etomoxir, an irreversible inhibitor of carnitine palmitoyl transferase 1a. Those results suggest that the FFAs released by SP16 treatment are utilized by the mitochondria through a process of β-oxidation.

**Fig. 3:**
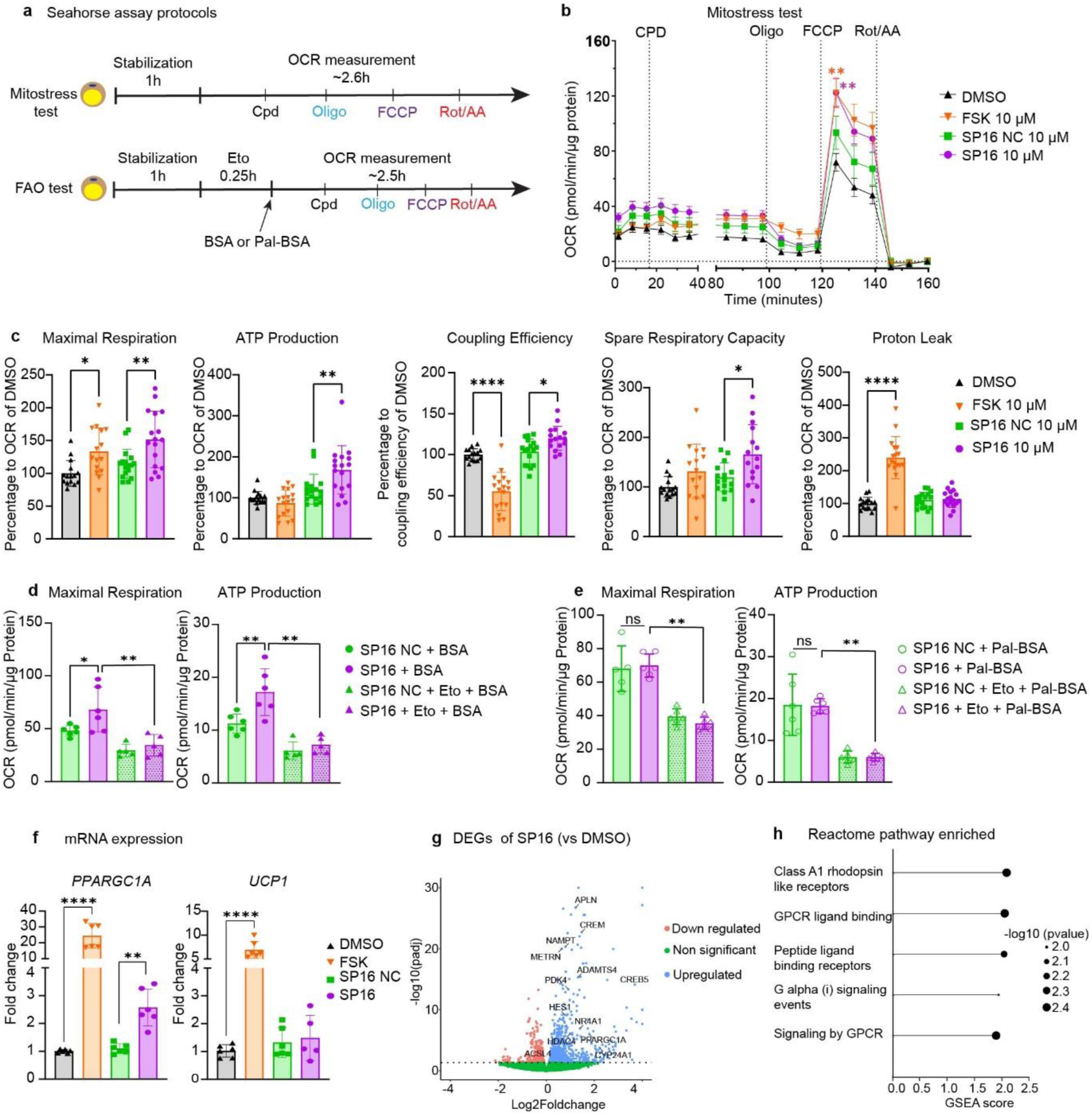
**SP16 increases β-oxidation and thermogenic markers in adipocytes**. Seahorse assay protocols (a), representative SP16-induced OCR of hMADS with mitostress test (b), SP16-induced quantitative maximal respiration, ATP production, coupling efficiency, spare respiratory capacity and proton leak of hMADS with mitostress test (c), SP16-induced quantitative maximal respiration and ATP production in hMADS pretreated with/without etomoxir in addition of BSA (d), ATP production in hMADS pretreated with/without etomoxir in addition of palmitate conjugated with BSA (e), mRNA expression of *PPARGC1A* and *UCP1* after 4 h treatment of FSK or SP16 (f), volcano plot of expression changes in SP16 vs DMSO. Labelled genes are involved in lipolysis, fatty acid activation, mitochondrial respiration and browning process (g). Pathway analysis of dysregulated genes of SP16 vs DMSO using Reactome database (GSEA) (h). Data are shown as mean ± SEM, n=6-18, *p < 0.05, **p<0.01, ****p<0.0001 by two-way ANOVA with Tukey’s post-hoc test (for b, d, e, f) or one-way ANOVA with Dunnett’s post-hoc test (for c). OCR: oxygen consumption rate, FAO: fatty acid oxidation, Oligo: oligomycin, FCCP: carbonyl cyanide-p-trifluoromethoxyphenylhydrazone, Rot/AA: rotenone/antimycin A, FSK: forskolin, SP16 NC: SP16 negative control, BSA: bovine serum albumin, Pal-BSA: palmitate conjugated with BSA, Eto: etomoxir, DEGs: differentially expressed genes.

### SP16 treatment modulates gene markers associated to lipolysis

To investigate the transcriptional response to SP16, we analyzed mRNA expression in hMADS cells following a 4 h treatment with 10 µM SP16. This analysis revealed a significant upregulation of *PPARG1A* expression. This finding aligns with the higher cAMP levels, which triggers PKA activation and subsequent downstream pathways, leading to increased expression of PGC-1α. PGC-1α, a transcriptional factor that plays a crucial role in orchestrating mitochondrial metabolism and activity, is known to regulate genes involved in mitochondrial biogenesis and dynamics^25^. However, treatment with SP16 does not induce the upregulation key genes associated with mitochondrial biogenesis (*TFAM, NRF1*) and genes involved in mitochondrial dynamics (*MFN1, MFN2*, *DRP1, OPA1, FIS1*) (Supplementary Fig. 2e - 2k). To further investigate the transcriptomic signature regulated by SP16, we conducted RNA sequencing on hMADS cells treated with SP16 for 4 h. We observed that SP16 treatment leads to increased expression of 950 genes and decreased expression of 689 genes (padj ≤ 0.05, FDR) (Fig. 3g). Specifically, we found increased expression of genes involved in regulation of lipolysis (*PDK4, NAMPT, CREM, CREB5*), fatty acid activation (ACSL4), mitochondrial respiration (*NR4A1, NR4A3*) and browning process (*APLN, ADAMTS4, HES1, PPARGC1A, HDAC4, CYP24A1, METRNL*) when adipocytes were treated with SP16^26,27^ (Fig. 3g). Furthermore, pathway enrichment analysis using the Reactome database (https://curator.reactome.org/PathwayBrowser) revealed that five pathways are enriched (Fig. 3h). Although no metabolic pathway is enriched, we observe a significant enrichment in pathways involved in G-protein-coupled receptor (GPCR) signaling. Altogether, these data show that the transcriptomic signature associated with SP16 treatment reflects the expected lipolysis-induced gene regulation. Additionally, the pathway analysis data suggests that the lipolytic phenotype may be mediated via GPCR signaling.

### SP16-induced lipolysis in adipocytes is regulated through GIPR

Based on the enrichment of GPCR pathways from RNA sequencing, we hypothesized that SP16 induces lipolysis in adipocytes through a GPCR. One GPCR known to induce lipolysis in adipocytes is the β3-adrenergic receptor^28^. To test the hypothesis that β3-adrenergic receptor mediates the effect of SP16, we antagonized β-adrenergic receptor with propranolol, a non-selective β-blocker. The lipolytic phenotype induced by SP16 is not blocked by propranolol (Supplementary Fig. 3a - 3b). Additionally, an additive lipolytic effect induced by SP16 and isoprenaline was observed (Supplementary Fig. 3c). Therefore, it can be concluded that SP16 does not induce lipolysis through β3-adrenergic receptor. Another known GPCR able to induce lipolysis upon activation is the GIPR^7,29^. We decided to evaluate the effect of SP16 on GIPR, utilizing CHO CRE-Luc2P cells overexpressing human GIPR (CRE-Luc2P hGIPR). SP16 induces cAMP levels in CRE-Luc2P hGIPR cells with an EC50 of 65 nM, demonstrating lower potency compared to the native ligand hGIP (EC50 = 18 pM) (Fig. 4a, 4m). To confirm the specificity for hGIPR, we introduced a GIPR antagonist^30^ prior to receptor activation using an EC80 agonist concentration (200 pM hGIP and 270 nM SP16, respectively). The GIPR antagonist inhibits cAMP activation in CRE-Luc2P hGIPR with an IC50 of 14 nM when activated with hGIP and an IC50 of 1.2 nM when activated with SP16 (Fig. 4b, 4n). Moreover, no cAMP induction by SP16 was observed on CRE-Luc2P empty cells (no GIPR receptor overexpression), confirming the specific activation of SP16 toward hGIPR (Supplementary Fig. 3d). To further investigate the interaction between SP16/hGIP and a GIPR antagonist on GIPR, we proceeded to a Schild regression analysis^31^. A rightward shift and no reduction in maximum response (Cmax) for both, SP16 (Fig. 4c) and hGIP (Supplementary Fig. 3f), are observed. This result indicated that SP16, similarly to hGIP, competes with the GIPR antagonist for binding to GIPR (pA2 = 8.7 when stimulated with SP16 and pA2 = 9.3 when stimulated with hGIP). pA2 value represents the potency of competitive reversible antagonism^31^. Radioligand binding result further confirmed that SP16 replaces ^125^I- GIP in binding to GIPR with a Ki value of 610 nM, whereas Ki value of hGIP is 536 pM (Fig. 4d and Supplementary Fig. 3e).

**Fig. 4:**
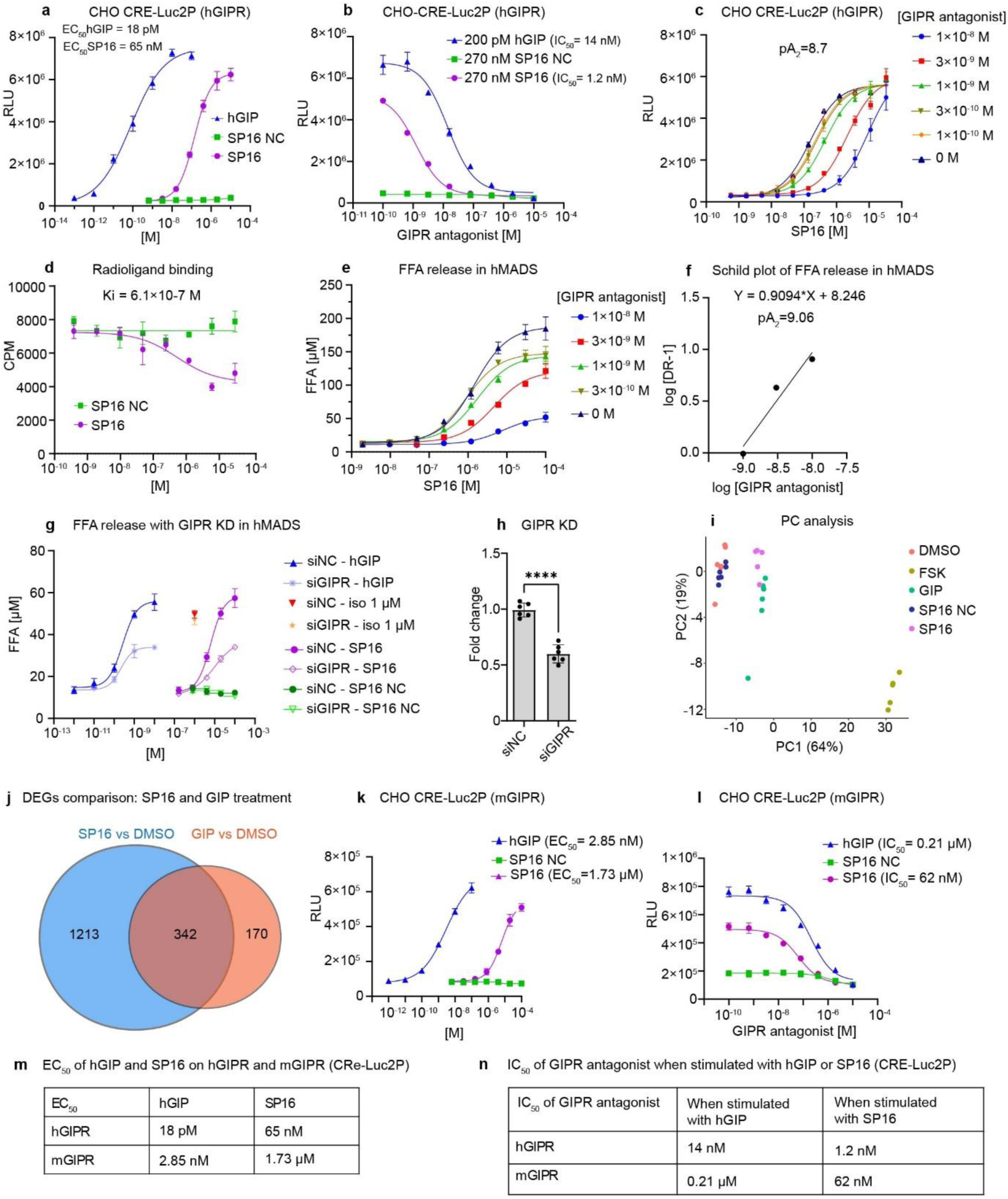
SP16-induced lipolysis in hMADS is regulated through GIPR. Luciferase activity induced by SP16 in CHO-CRE-Luc2P cells overexpressing hGIPR (a), SP16-induced luciferase activity of CHO CRE-Luc2P cells overexpressing hGIPR pre-treated with GIPR antagonist (b), SP16 induced luciferase activity of CHO CRE-Luc2P cells overexpressing hGIPR pre-treated with different concentrations of GIPR antagonist (c), competitive binding of SP16 to hGIPR membrane vs ^125^I-GIP (d), SP16-induced FFA release in hMADS when treated with GIPR antagonist and SP16 (e), competitive antagonist analysis (Schild plot) obtained from FFA release in hMADS (f), FFA release in cells transfected with siRNA targeting GIPR or negative control siRNA (g), fold change in GIPR gene expression in hMADS transfected with siRNA targeting GIPR or negative control siRNA (h), PCA analysis based on top 500 variance genes (i) . Overlap of DEGs between SP16 vs DMSO and GIP vs DMSO contrasts (adjp <0.05, FDR) (i) SP16-induced luciferase activity of CHO CRE-Luc2P overexpressing mGIPR (k), and pre- treated with GIPR antagonist (l), summarized EC50 of hGIP and SP16 on CRE-Luc2P cells overexpressing hGIPR or mGIPR (m), summarized IC50 of GIPR antagonist when CRE-Luc2P cells overexpressing hGIPR or mGIPR cells were stimulated with hGIP or SP16 (n). Data are shown as mean ± SEM, n=4-8, ****p < 0.0001 by 2-tailed unpaired t-test. hGIPR: human GIPR, mGIPR: mouse GIPR, RLU: relative light unit, DR: dose ratio, siGIPR: siRNA targeting GIPR, siNC: siRNA negative control, KD: knockdown, FFA: free fatty acid, SP16: SP16 negative control, iso: isoprenaline.

Next, we assessed whether GIPR antagonism or siRNA-mediated knockdown (KD) attenuates lipolysis induced by SP16. Indeed, GIPR antagonism suppresses the FFA release driven by SP16. Schild regression analysis on hMADS lipolysis induced by SP16 revealed that the GIPR antagonist had a pA2 value of 9.06 (Fig. 4e, 4f). A similar pA2 value of the GIPR antagonist was obtained when treating hMADS cells with hGIP and the GIPR antagonist (Supplementary Fig. 3g, 3h). Additionally, FFA release induced by hGIP or SP16 is significantly reduced in hMADS cells treated with GIPR siRNA compared to cells transfected with negative control siRNA (siNC) (Fig. 4g). The knockdown efficiency of GIPR was approximately 40% (Fig. 4h). Lipolysis induced by 1 µM isoprenaline remains unchanged when cells were treated with GIPR antagonist.

Moreover, RNA sequencing of hMADS cells treated with DMSO, FSK, SP16 NC, SP16, or GIP revealed three distinct clusters in principal component analysis (PCA): cluster 1 (DMSO and SP16 NC), cluster 2 (SP16 and GIP), and cluster 3 (FSK) (Fig. 4i). This clustering indicates that SP16 and GIP elicit a partly similar transcriptomic signatures in adipocytes. Specifically, 342 differentially expressed genes (DEGs) are common between the SP16 and GIP treatment groups, while 1213 DEGs are uniquely regulated by SP16 and 170 by GIP (Fig. 4j). Notably, genes involved in lipolysis, fatty acid activation, mitochondrial respiration, and the browning process, which are modulated by SP16 (Fig. 3g), are also higher in the GIP-treated group (Supplementary Fig. 3i). Interestingly, SP16 also seems to regulate genes independently of GIPR, suggesting potential interactions with other binding partners. To investigate the translatability of SP16-induced cAMP production from human to mouse cells, we utilized CRE- Luc2P reporter cells overexpressing mouse GIPR (mGIPR). Both hGIP and SP16 activate mGIPR with EC50 values of 2.85 nM and 1.73 µM, respectively (Fig. 4k). The activation of both hGIP and SP16 is suppressed by the introduction of the GIPR antagonist^30^ with IC50 values towards mGIPR of 210 nM and 62 nM for hGIP and SP16, respectively (Fig. 4l). This finding suggests that SP16 and hGIP remain potent on the mGIPR despite a difference of potency compared to the hGIPR.

In summary, our findings demonstrate a conserved binding and activation of the GIPR by SP16. Furthermore, we provide evidence that the lipolytic activity of SP16 is predominantly mediated through its interaction with GIPR. Transcriptomic analyses revealed that the gene expression profile induced by SP16 is partially overlapping with canonical GIPR signaling, while also exhibiting distinct features unique to SP16. This duality suggests that SP16 may exert its biological effects through both GIPR-dependent and GIPR-independent mechanisms.

### SP16 binds to the insulin receptor and inhibits insulin receptor signaling pathways

In order to identify additional molecular targets of SP16, a binding and functional panel consisting of 87 pharmacologically relevant receptors and proteins of interest^32^ was performed (Supplementary Table 5, 6). SP16 shows 69.3% inhibition of control specific binding to insulin receptor (INSR) kinase and 79.9% inhibition to acetylcholinesterase (AChE) (Fig. 5a). Despite the higher percentage inhibition of AChE, INSR is well-known for its crucial roles in growth, proliferation, and metabolism, making it relevant to investigate the relationship between SP16 and INSR^33^. To assess the interaction between SP16 and INSR, a surface plasmon resonance (SPR) binding assay was employed. Biotinylated SP16 and biotinylated SP34 (SP16 scrambled) were immobilized on the chip surface, and recombinant INSR protein was injected over the peptide coupled sensor. The results indicated that SP16 binds to INSR with a KD in the range of 185 nM, suggesting a good binding affinity between SP16 and the INSR (Fig. 5b, 5c). Conversely, the insulin receptor does not exhibit binding to SP34 (Supplementary Fig. 4a, 4b), confirming the direct interaction between SP16 and INSR.

**Fig. 5:**
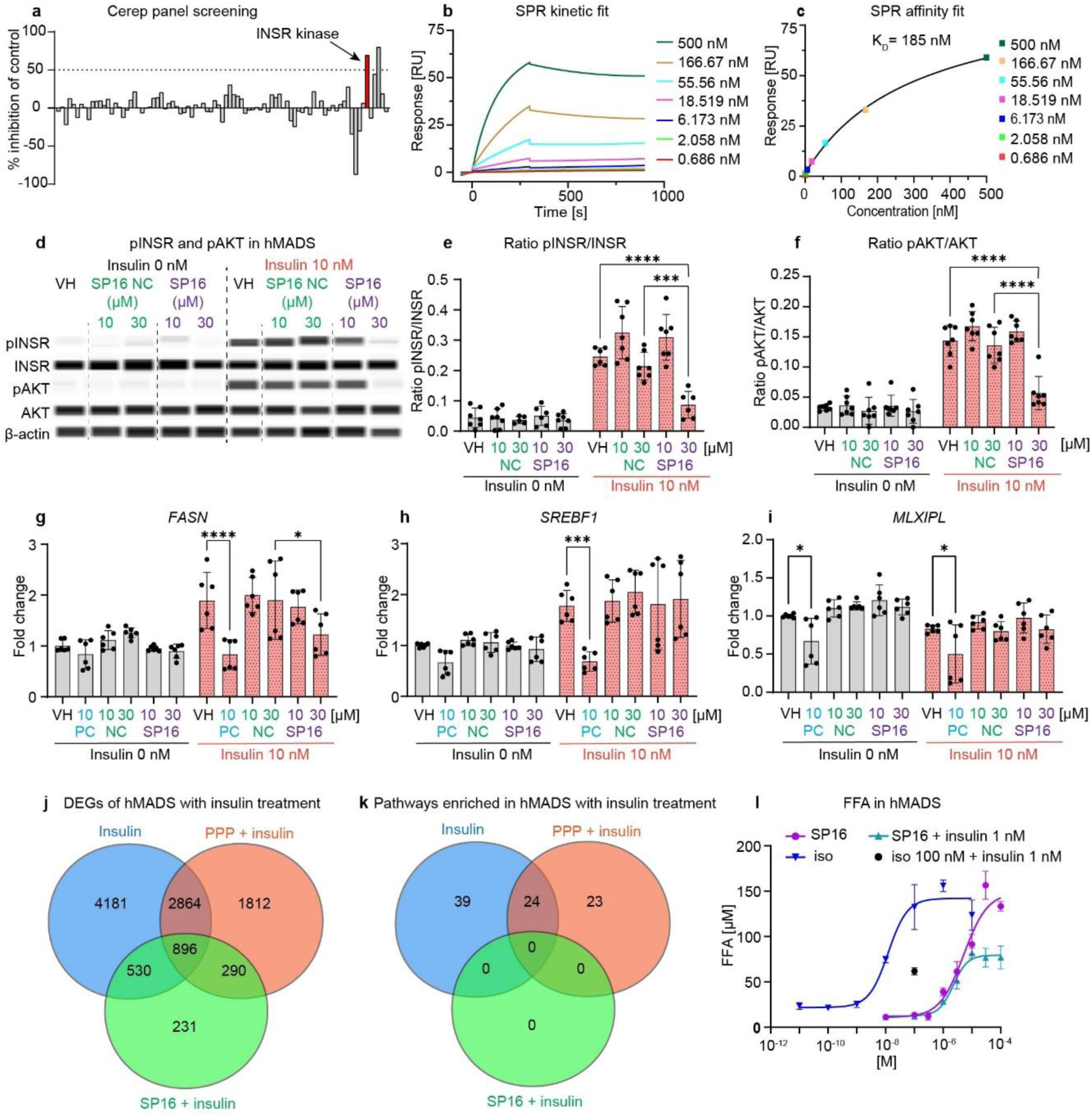
SP16 binds to the insulin receptor and modulates insulin receptor signaling pathway. Binding of SP16 in the Cerep panel 87-assay screen (a), SPR binding assay of INSR to biotinylated SP16: kinetic fit (b), affinity fit of to INSR (c), SP16 modulates quantification of pINSR and pAKT in hMADS shown by Western blot in presence or absence of 10 nM insulin (d), pINSR/INSR ratio (e), pAKT/AKT ratio (f), mRNA expression changes by SP16 in presence or absence of 10 nM insulin: *FASN* (b), *SREBF1* (h), *MLXIPL* (i), FFA release in hMADS induced by SP16 in presence of 1 nM insulin (j). DEGs (adjp <0.05) from following contrasts, Insulin vs DMSO, PPP + insulin vs DMSO + insulin and SP16 + insulin vs DMSP + insulin (j). Pathway analysis of dysregulated genes of above-mentioned contrasts Reactome database (GSEA) (h) Data are shown as mean ± SEM, n=6-8, *p < 0.05, **p<0.01, ***p<0.01, ****p<0.0001 by two-way ANOVA with Tukey’s post-hoc test. INSR: insulin receptor, AChE: acetylcholinesterase, RU: response unit, SPR: surface plasmon resonance, SP16 NC: SP16 negative control, NC: SP16 NC, VH: vehicle, PC: positive control (PPP - Picropodophyllin), FFA: free fatty acid.

The impact of SP16 on the insulin signaling pathway was evaluated. Under basal conditions, SP16 does not modulate the phosphorylation of AKT in hMADS (Fig. 5b and Supplementary Fig. 4c). However, upon stimulation with 10 nM insulin, SP16 at a concentration of 30 µM significantly inhibits approximately 70% of pINSR and pAKT compared to the SP16 NC, while SP16 at 10 µM does not exhibit any inhibition of pINSR and pAKT (Fig. 5d, 5e, 5f). When the concentration of insulin increased to 100 nM, the inhibitory effect of SP16 is diminished (Supplementary Fig. 4d, 4e, 4f). As insulin signaling pathway is known to play a crucial role in regulating lipogenesis^33,34^, we examined the expression of lipogenic markers in hMADS following treatment with insulin and SP16, including *FASN, SREBP-1c, MLXIPL, USF1,* and *USF2*. Picropodophyllin (PPP), an insulin like growth factor 1 receptor inhibitor, was used as a positive control. The results showed that 24 h insulin treatment upregulates the expression of *FASN* and *SREBF1* (Fig. 5g, 5h). When treated with SP16 or PPP in the presence of 10 nM insulin, the only expression of *FASN* is significantly lower (Fig. 5g). The expression of other lipogenic markers (*MLXIPL, USF1, USF2*) remains unchanged (Fig. 5h, Supplementary Fig. 4g, 4h).

To determine whether the inhibition of INSR by SP16 modulates the transcriptional signature regulated by insulin signaling, we conducted RNA sequencing on hMADS cells treated with insulin, PPP and insulin, and SP16 and insulin. Our results showed that in the presence of insulin, SP16 significantly regulates 1947 genes, with 1426 genes overlapping with those regulated by insulin, and 1186 genes overlapping with those regulated by PPP and insulin (Fig. 5j, adjp < 0.05, FDR). Detailed information on DEGs from each treatment is depicted in Supplementary Fig. 5a, 5b, 5c. Pathway analysis using the Reactome database revealed that insulin treatment in hMADS cells modulated 63 pathways (Fig. 5k), with 51 pathways positively enriched and 12 pathways negatively enriched (Supplementary Fig. 5d). Treatment with PPP reversed the regulation of 24 pathways modulated by insulin (Fig. 5k, Supplementary Fig. 5d). In contrast, no pathways are significantly enriched when cells were treated with SP16 and insulin, indicating no overlapping pathways between SP16 and either insulin or PPP (Fig. 5j, Supplementary Fig. 5d). Insulin is well described for its anti-lipolytic effect through the stimulation of cAMP degradation via the activation of phosphodiesterase 3B (PDE-3B)^35^. Therefore, we assessed the inhibitory effect of SP16 on the anti-lipolytic effect of insulin. The results showed that insulin suppressed FFA release similarly regardless of the stimulus, SP16 or isoprenaline (Fig. 5l), suggesting that the inhibition of INSR by SP16 does not lead to increase of FFA. In conclusion, SP16 binds INSR and inhibits insulin pathway in hMADS cells at a concentration of 30 µM. Although this inhibition by SP16 modulates gene expression upon insulin stimulation, it does not significantly interfere with the metabolic pathways regulated by insulin.

### SP16 induces lipolysis and improves glucose tolerance in DIO mice

It is well described that activation of the GIPR improves metabolic parameters in mice fed with HFD^7,11,36^. Based on this, we hypothesized that SP16, which we have identified as a GIPR agonist, may exert beneficial effects on metabolic regulation *in vivo*. Firstly, the pharmacokinetic (PK) profile of SP16 was evaluated via intraperitoneal (i.p.) administration in mice at a dose of 300 nmol/kg. However, 15 minutes after i.p. injection, SP16 was not detectable in plasma. This rapid elimination from plasma indicates a short half-life of SP16. Therefore, we performed acute treatment to assess the metabolic effects of SP16 in mice considering its short half-life with two doses, 25 mg/kg and 75 mg/kg (experimental protocol depicted in Fig. 6a). GIPR agonist (GIPRA) (patent number: WO/2018/181864) was used as positive control.

**Fig. 6:**
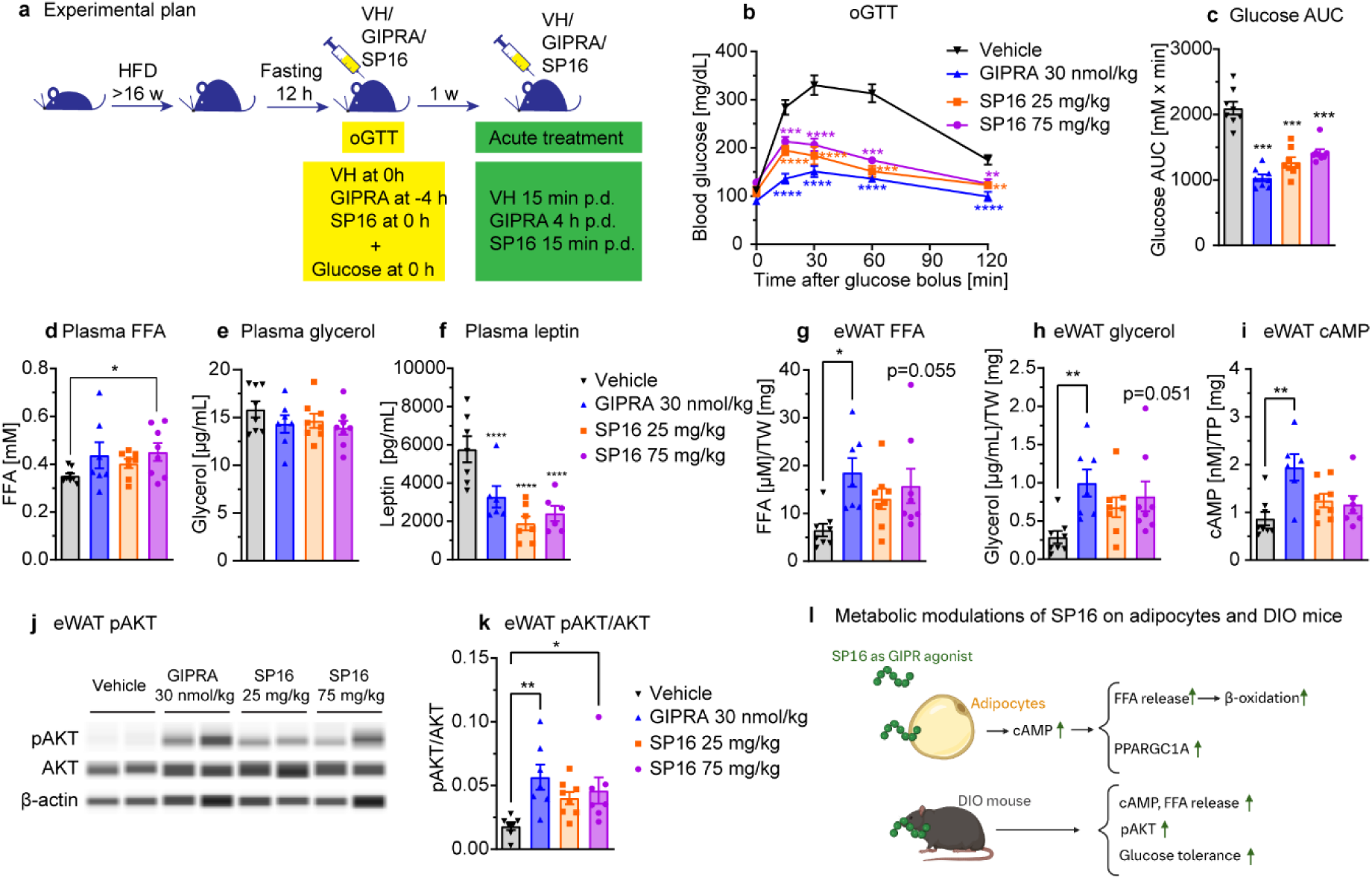
**SP16 improves lipid and glucose metabolism in DIO mice**. Experimental protocol (a), effect of SP16 on blood glucose during oGTT (b), glucose area under the curve during the oGTT period (c), plasma FFA (d), plasma glycerol (e), plasma leptin (f), normalized FFA release from eWAT (g), glycerol release from eWAT (h), cAMP concentration in eWAT (i), pATK level in eWAT (j), pAKT/AKT ratio in eWAT (k) during GIPRA and SP16 treatment, metabolic modulation of SP16 as a GIPR agonist on adipocytes and DIO mice (l). Data are mean ± SEM, n=7-8, *p < 0.05, **p<0.01, ***p<0.01, ****p < 0.0001 by two-way ANOVA with Tukey’s post-hoc test (for d) and one-way ANOVA with Dunnett’s post- hoc test (for e, f, g, h, I, j, k, m). GIPRA: GIPR agonist, HFD: high fat diet, VH: vehicle, p.d. post dosing, oGTT: oral glucose tolerance test, AUC: area under the curve.

In the oral glucose tolerance test (oGTT), SP16 significantly improves glucose tolerance in DIO mice (Fig. 6b), with a significant decrease in glucose area under the curve (AUC) of 40% (Fig. 6c). After 60 min of glucose load, SP16 exhibits a similar improvement in glucose tolerance compared to the GIPRA. Additionally, a 15-min treatment with SP16 resulted in higher level of plasma FFA and lower leptin level compared to vehicle treated mice (Fig. 6d, 6e, 6f). Different adipose tissue depots (iWAT, eWAT, iBAT) and the liver were investigated *ex vivo* regarding changes in lipolysis, cAMP levels and pAKT. A trend for elevated levels of FFA, glycerol, and cAMP (Fig. 6g, 6h, 6i), as well as a significant increase in pAKT in eWAT was observed (Fig. 6j, 6k). However, SP16 failed to significantly affect FFA, glycerol, cAMP and pAKT in iWAT, iBAT, and the liver at the tested dose (Supplementary Fig. 6).

In conclusion, acute treatment with SP16 on mice fed with HFD significantly improves glucose tolerance, increases pAKT levels in eWAT, and reduces plasma leptin levels (Fig. 6l), suggesting that the GIPR activity of SP16 seen *in vitro* translates to *in vivo*.

## Discussion

The rising incidence of obesity globally presents a significant challenge to healthcare systems and underscores the urgent need for effective therapeutics to support patients living with obesity. Enhancing adipocyte metabolism to promote thermogenesis and EE offers an avenue to improve existing therapies for obesity and its associated metabolic disorders.

In this study, we confirmed the association of AAT with obesity by demonstrating a significant downregulation of SERPINA1E at both mRNA and protein level of DIO mice compared to lean controls. This observation is consistent with the previous reports showing reduced SERPINA1E expression in other obesity models, including ob/ob mice and in db/db mice, while other members of the SERPINA1 family did not exhibit similar patterns^13,37^. Unlike other SERPINA1 isoforms, the protease target of SERPINA1E has not yet been identified (according to Uniprot.org). Therefore, it is interesting to further investigate the functional implications of SERPINA1E in obesity and related metabolic diseases.

Importantly, we reported for the first time that SP16, a synthetic peptide derived from obesity-associated protein AAT, activates catabolic processes in adipocyte through induction of lipolysis. Mechanistically, we showed that SP16 exerts its effects through activation of the GIPR, leading to increased FFA release and β-oxidation in hMADS. Furthermore, SP16 administration improves metabolic parameters in DIO mice. Our findings align with a study by Okgawa *et al*., whose work demonstrated the beneficial effect of hepatic AAT in enhancing adaptive thermogenesis, EE and glucose tolerance^24^. The underlying mechanism is suggested to involve the interplay between AAT and Eph receptor B2 (EphB2)^24^. EphB2 was not assessed in our study and is not included in binding and functional panel. Further experiments would be necessary to evaluate the interaction between EphB2 and SP16, since EphB2 is expressed in hMADS (Supplementary Fig. 7a)

The results obtained from binding and functional panel^32^, which comprises 87 molecular targets, revealed that SP16 exhibits inhibitory activity against AChE. AChE hydrolyzes acetylcholin (ACh) within the central and autonomic nervous systems, as well as at neuromuscular synapses^38^. Therefore, the inhibition of AChE leads to increase ACh levels and improve cognitive function therapeutically^39^. However, it is important to note that elevated ACh levels can potentially lead to a cholinergic crisis^39^. Moreover, SP16 has been demonstrated to traverse an artificial 3D blood-brain barrier (BBB) system and the BBB in mice following intranasal delivery^40^. Although our study did not further investigate the effect of SP16 on AChE, it is crucial to carefully consider the implications of AChE inhibition, e.g. in DIO mice, since AChE is expressed in adipocytes (Supplementary Fig. 7a).

Despite few receptor binding activities identified using the binding and functional panel^32^, literature suggests that SP16 has several binding partners. Toldo *et al*. has described SP16 as a potential partner for LRP1 agonist^18^. We could confirm those finding using SPR (Supplementary Fig. 7b-7e). The LRP1 receptor is a scavenging receptor that plays a critical role in the regulation of energy homeostasis, lipid metabolism, and food intake^23,41^. Hepatic LRP1 knockout mice exhibit enhanced insulin resistance, HFD-induced obesity, dyslipidemia, and hepatosteatosis^41^. Similarly, GABAergic neuron-specific LRP1-deficient mice demonstrate increase in food intake and decrease in EE, locomotor activity, insulin resistance, and glucose intolerance^42^. Therefore, we investigated the involvement of LRP1 receptor in the lipolytic effect of SP16 in hMADS using an LRP antagonist (RAP) and genetic knockdown of the LRP1 with siRNA. However, our data suggested that in hMADS model, SP16 does not induce lipolysis through LRP1 (Supplementary Fig. 7f - 7i) despite a high expression of LRP1 and other members of the lipoprotein family (Supplementary Fig. 7a).

We demonstrated that in hMADS, at a concentration of 30 µM, SP16 antagonizes the INSR and its signaling pathway. The INSR inhibition might translate *in vivo* by an impairment of the uptake of glucose from the bloodstream into tissues, resulting in elevated blood glucose levels and a compensatory response from the pancreas such as increased insulin production^43^. Consequently, the implications of INSR inhibition, particularly its role in the pathogenesis of insulin resistance and related metabolic disorders, require careful consideration. Interestingly, SP16 still shows a significant improvement in glucose tolerance in DIO mice. We observed that *in vitro* INSR inhibition by SP16 on hMADS does not translate into mouse adipocytes, demonstrated by the less pronounced inhibition of pINSR and pAKT in 3T3-L1 cells (Supplementary Fig. 8a, 8b). Furthermore, as widely described, it is expected that DIO mice may already exhibit insulin resistance, which could explain the less prominent inhibitory effect of SP16 on INSR in these mice^44,45^.

In this study, we demonstrated that SP16 functions as a GIPR agonist, with an EC50 value of 65 nM which is less potent compared to the endogenous hGIP (EC50 = 18 pM) in the CRE- Luc2P hGIPR system (Fig. 4m). On hMADS cells with FFA release read-out, both SP16 (EC50 = 7.43 µM) and hGIP (EC50 = 1.1 nM) exhibited lower potency compared to CRE-Luc2P hGIPR cells. This could be attributed to differences in receptor density between cells or differences in the experimental read-out between the two cell models. Notably, at physiological condition, the plasma concentration of AAT is relatively high (1-2 g/L or 19-38 µM), which make the EC50 of SP16 in the lipolysis assay meaningful. Furthermore, both SP16 and hGIP, showed reduced potency on mGIPR compared to hGIPR. Research findings suggest that although both GIP receptors have similar agonist affinities and potencies for Gαs activation in response to endogenous hGIP, hGIPR is more susceptible to desensitization than mGIPR^46^, thus providing an explanation for the observed differences.

Previous studies have reported an inverse correlation between the expression of GIPR in adipose tissue and insulin resistance^47^. It has been shown that treatment with GIP lowered insulin-induced pAKT in adipocytes differentiated from adipose stem cells of obese patients compared to lean individuals^47^. Similarly, we have observed that SP16, as a GIP mimic peptide, lowers the phosphorylation level of INSR and AKT under stimulation of insulin.

Endogenous hGIP has a short half-life of approximately 7 minutes in healthy individuals due to enzymatic degradation by dipeptidyl peptidase 4^48^. Similarly, our PK study in mice revealed that SP16 is quickly eliminated from plasma. Given its low molecular weight (2.2 kDa), SP16 is likely to be rapidly cleared through renal elimination. Additionally, SP16 may be susceptible to cleavage by proteases at the injection site or in plasma. Several strategies, such as the use of poloxamer and encapsulation of SP16 within poly(lactide-co-glycolide) particles for extended release, were explored to improve the half-life of SP16^49^. However, PK profiles of these formulations did not show any improvement (data not shown). Extending half-life of SP16 through chemical structure modification can be a promising method to improve PK profile of SP16. Nevertheless, in DIO mice, SP16 at doses of 25 mg/kg and 75 mg/kg exhibited metabolic benefits similar to a GIPR agonist. It would also be of interest to investigate in follow-up studies whether chronic treatments with SP16 exhibit beneficial effects.

In conclusion, we have demonstrated that the AAT-derived peptide SP16, acts as a novel GIPR agonist, and shows beneficial effects on adipocytes and metabolic functions in DIO mice. These results highlight SP16’s potential as a therapeutic candidate for obesity and related metabolic disorders. While further optimization is necessary to improve its pharmacokinetic properties, specificity, and potency, these challenges offer valuable starting points for future development.

## Material and Methods

### In silico peptide selection

To identify novel peptide associated with obesity, top 20 genes associated with body weight were selected based on their HuGE score (from Common Metabolic Diseases Knowledge Portal - https://hugeamp.org). Subsequently, the proteins encoded by the selected 20 genes were subjected to analysis using the Uniprot database. This analysis aimed to determine if any of the proteins were predicted to undergo cleavage, resulting in the generation of peptides. The resulting Uniprot-annotated peptides obtained from the previous step were compared with the peptidomic data obtained from a study conducted by Arapidi *et al*.^50^. The peptides which were found to be detected in the peptidome database, were selected for further analysis. For validation, the human genes (obtained after Uniprot-annotated peptide selection step) were compared with genes coding for proteins originating from the mouse peptidomes from the study of Madsen *et al*.^19^.

### Peptide synthesis

SPAAT, SP16, SP16 negative control containing 5 alanine residues (SP16 NC), SP34 (SP16 scrambled) (amino acid sequences described in Supplementary Table 2) were synthesized using microwave-assisted solid-phase peptide synthesis (SPPS) and standard Fmoc deprotection strategies. Peptides were purified to >90% peak purity at 214 nm as assessed by high performance liquid chromatography, and the molecular weights were confirmed by time-of-flight mass spectrometry. Peptides are N-terminally acetylated (“Ac-“) and C-terminally amidated (“-NH2). “Nle” refers to L-norleucine. All other amino acids are denoted by their one letter codes and are in their L-forms.

### Adipocyte culture and differentiation

Human multipotent adipose-derived stem cells (hMADS) were obtained following the protocol described elsewhere^51^. hMADS cells were cultured in DMEM low glucose (1 g/L) with L-GlutaMAX (Gibco, 11885084) supplemented with 10% fetal bovine serum (FBS) (Gibco, 11500064), 1% Pen/Strep (Gibco, 15070063), 15 mM HEPES (Gibco, 15630056) and 2.5 ng/ml bFGF-2 (Peprotech, 100-18B-50UG). Tissue culture-treated surfaces were used for seeding the hMADS cells. The cells were induced to differentiate at day 2 post confluence (designated as day 0) in Omental Adipocyte Differentiation Medium (Zenbio, OM-DM°500mL).

Medium was changed every other day. At day 7, the media were replaced with Omental Adipocyte Maintenance Medium (Zenbio, OM-AM°500mL). Starting from day 9, the cells contained visible lipid droplets and were used for further experiments. To obtain brown adipocytes, Omental Adipocyte Maintenance Medium was supplemented with 1 µM rosiglitazone (Sigma, R2408-10MG).

Primary human stromal vascular fractions (hSVF) were isolated following the established protocol^52^. The hSVF cells were cultured in DMEM low glucose (1 g/L) with L-GlutaMAX supplemented with 10% FBS, 1% Pen/Strep and 15 mM HEPES. The cells were then seeded in tissue culture treated surface. Adipogenesis was induced when the cells reached 100% confluency, similar to hMADS cells, using Omental Adipocyte Differentiation Medium and Omental Adipocyte Maintenance Medium from Zenbio. The cells were subjected to assays at day 11 of differentiation.

The 3T3-L1 cell line was purchased from ATCC and cultured in DMEM 4.5 g/L glucose (Gibco, 11965092) supplemented with 10% FBS, 1% Pen/Strep. When the cells reached confluency (designated day 0), the medium was changed to differentiation medium, which consisted of proliferation medium supplemented with 0.25 µM dexamethasone (Sigma, 265005-100 MG), 0.5 mM IBMX (Sigma, I5879-250 MG), 1 µM rosiglitazone (Merck, R2408-10 MG) and 1.67 µM insulin (Sigma, I9278-5 ML). On day 3, the medium was changed to maintenance medium which contained proliferation medium supplemented with 1.67 µM insulin. Starting from day 5, the medium was changed back to proliferation medium. Proliferation medium was changed every other day. The cells were used for assays at day 9 or 10 of differentiation.

### Cell treatment with insulin

The 9-day old differentiated hMADS cells were starved overnight with starvation medium consisting of DMEM without glucose supplementation with 0.5% BSA (37 °C, 5% CO2). On the following day, for western blot assays, cells were incubated with compounds for 15 min (37 °C, 5% CO2). Insulin was added into corresponding samples and incubated for another 15 min (37 °C, 5% CO2). When the incubation time was finished, cells were washed with PBS and lysed as previously described. For mRNA expression determination, cells were pre-incubated with compounds for 30 min. Insulin was added to the corresponding samples and incubated for another 24 h (37 °C, 5% CO2). When the incubation was finished, the cells were washed with PBS and lysed with RLT buffer supplemented with 1% 2-mercaptoethanol for mRNA isolation and RT-qPCR.

### Lipolysis assay

The lipolysis assay was performed using Krebs-Ringer bicarbonate buffer (KRBB, K4002- 10X1 L) supplemented with 2% bovine serum albumin (BSA).

For hMADS, hSVF, 3T3-L1, the cells were starved with lipolysis buffer in an incubator (37 °C, 5% CO2) for 2 h. The buffer was then replaced with compounds diluted in lipolysis buffer and incubated for an additional 3 h. After the incubation period, the supernatant was collected, and the concentration of free fatty acid (FFA) was determined using a non-esterified fatty acid (NEFA) kit (WAKO, 434-91795, 436-91995) following the manufacturer’s protocol. The absorbance was measured at a wavelength of 546 nm using Spectramax 384 Plus (Molecular Device), and the FFA concentration was calculated based on the NEFA standard concentration.

For lipolysis with adipose tissue, inguinal white adipose tissue (iWAT), epididymal WAT (eWAT) and interscapular BAT (iBAT) obtained from mice were cut into small pieces (approximately 10 mg/piece for iWAT and eWAT and 5 mg/piece for BAT) and placed into a 96-well plate containing pre-warmed lipolysis buffer. The plate was incubated for 1 h at 37 °C, 5% CO2 as a stabilization step, and the adipose tissue pieces were transferred to a plate containing the desired compounds diluted in lipolysis buffer. The plate was incubated at 37 °C, 5% CO2 for 3 h. The supernatant was collected, and the FFA concentration was determined as described above. The FFA concentration was then normalized to the weight of each tissue piece.

### cAMP assay

The differentiated hMADS cells were stimulated with compounds for 30 min (37 °C, 5% CO2) in KRBB supplemented with 250 µM IBMX. Cells were washed with PBS and homogenized in 0.1 N HCl (Enzo Life Sciences, ADI-900-067A) supplemented with 0.1% Triton X-100 (Merck, T8787-50 ML) for 10 min, and were then centrifuged at 1000 × g for 5 min. 100 µL of cell lysate was transferred for the ELISA assay with acetylated format following manufacturer’s protocol (Enzo Life Sciences, ADI-900-067A). The cAMP levels were determined via absorbance measurement at 410 nm with a SpectraMax 384 Plus.

### Measurement of oxygen consumption

Oxygen consumption rate (OCR) was measured using Seahorse XF96 Pro Analyzer (Agilent). For the acute mitochondrial stress test, the assay media consisted of XF DMEM Medium pH 7.4 supplemented with 20 mM Glucose (Agilent), 2 mM L-Glutamine (Agilent), 1 mM Pyruvat (Agilent). The 9-day old differentiated hMADS were incubated with assay media in a non-CO2 incubator at 37°C for 1 h. Seahorse sensor cartridges were hydrated the night before and the ports were loaded to achieve the following final concentrations after injection: port A (compound), port B (oligomycin 1.2 µM), port C (FCCP, 2 μM), port D (rotenone 1 μM and antimycin A 1 μM). Oligomycin, FCCP, rotenone/ antimycin A were purchased from Agilent as in mitochondrial stress test kit (103015-100).

For the long-chain fatty acid oxidation (FAO) test, the Seahorse XF Palmitate Oxidation Stress Test Kit and FAO Substrate (Agilent, 103693-100, 102720-100) were utilized. The 9-day old differentiated hMADS cells were incubated overnight (37 °C, 5% CO2) with substrate limited growth media (DMEM Medium pH 7.4 supplemented with 0.5 mM Glucose, 1 mM L-Glutamine, 1% FBS and 0.5 mM L-Carnitine). On the following day, the cells were first incubated for 1 h in non-CO2 incubator with substrate-limited assay media. Etomoxir 100 µM (Merck, 236020-5MG) was added to the corresponding wells. After 15 minutes, palmitate conjugated with BSA or BSA was added, and the cells were immediately subjected to Seahorse measurement. The measurement setup is described in detail in Supplementary Table 3.

After the measurement, cells were washed with PBS (Sigma, D8537-500ML) and lysed with RIPA buffer (Sigma, R0278-500ML) supplemented with Halt^TM^ Protease and Phosphatase inhibitor Cocktail 100x (Thermo Fisher, 78440). The total protein concentration was determined using Pierce^TM^ BCA Protein Assay Kits (Thermo Fisher, 23227) following manufacturer’s protocol. The absorbance was measured at a wavelength of 562 nm using the SpectraMax 384 Plus. Total protein concentration was calculated based on BSA standard and was used for OCR normalization.

### RNA isolation and cDNA synthesis

Cells were lysed with RLT buffer (Qiagen, 79216) supplemented with 1% 2-mercaptoethanol (Sigma, M6250-10ML). RNA was extracted utilizing RNeasy kit (Qiagen, 74104) following the manufacturer’s protocol. RNA concentration was measured using Nanodrop spectrophotometer (Implen). cDNA was synthesized from 750 ng of total RNA in 30 µL reaction using the High-Capacity cDNA Reverse Transcription Kit (Thermo Fisher, 4368814).

### Real time quantitative PCR

Relative mRNA was quantified using Taqman primers purchased from Thermo Fisher (Supplementary Table 4). cDNA was diluted 1:5 and RT-qPCR was performed with TaqMan^TM^ 2x Fast Advanced Master Mix (Applied Biosystems, 4444558) in QuantStudio 6 or 7 Flex Real Time (Applied Biosystems). Cycles for genes of interest were normalized on the housekeeping gene TBP.

### Immunoblotting with Sallysue

Protein levels were normalized to 250 ng/mL. Reagents used for western blotting with Sallysue, including antibody diluent, streptavidin, goat anti-mouse secondary antibody, goat anti-rabbit secondary antibody, capillaries and sample plates were all obtained from Bio- techne (SM-W001, DM-001, DM-002). Samples were prepared according to the recommended manufacturer’s protocol. Primary antibodies were obtained from Cell Signaling Technology, including insulin receptor β (4B8) rabbit mAb (3025S), phospho-IGF-I receptor β (Tyr1135/1136)/insulin receptor β (Tyr1150/1151) (19H7) rabbit mAb (3024S), AKT (pan) (E7J2C) mouse mAb (58295S), phospho-AKT antibody (9271L), β-Actin antibody (4967L). The chemiluminescent signal of primary antibodies was detected and quantified using the software Compass for SW (ProteinSimple). Phosphorylation levels of insulin receptor and AKT were normalized to protein levels of insulin receptor and AKT, respectively.

### Gene knockdown with siRNA

Transfection of hMADS cells was performed at day 4 of differentiation using Lipofectamine RNAiMAX Reagent (Invitrogen, 13778075), following the manufacturer’s instructions. The siRNA concentrations used for LRP1 and GIPR knockdown were 6 nM and 20 nM, respectively. ON-TARGETplus SMARTpool human LRP1 siRNA (L-004721-00-0010), ON-TARGETplus SMARTpool human GIPR siRNA (L-005514-00-0010), ON-TARGETplus Control Pool Non-Targeting pool siRNA (D-001810-10-20) were purchased from Dharmacon. On day 7, a second transfection was carried out on the cells using the same conditions as the first one. On day 9, a portion of the cells was subjected to the lipolysis assay following previously described methods, while another portion of the cells was lysed for the RT-qPCR assay.

### Cre-Luc2P reporter assay

The study utilized CRE-Luc2P reporter cell lines without receptor overexpression and CRE- Luc2P reporter cell lines overexpressing recombinant receptors, including human GIPR and mouse GIPR. These cell lines were generated following established protocols^53^. Details of the culture media for each reporter cell line can be found in the Supplementary Table 7. Sequences of hGIP is described in Supplementary Table 2. On the day before, the cells were seeded at a concentration of 30,000 cells/well on the opaque CulturPlate-384 white (PerkinElmer, 6007680). The following day, cells were treated with peptides in KRBB supplemented with 10 mM HEPES buffer and 0.5% BSA for 4 h (37 °C, 5% CO2). After the incubation time, the buffer was replaced with 15 µL Bright-Glo Luciferase assay reagent (Promega, E2650) and incubated at room temperature with shaking for 10 min. Subsequently, 15 µL of PBS was added to each well and luminescence levels were measured with EnVision Multilabel Plate Reader (PerkinElmer).

### Functional and binding panel 87 assays

Assay categories in the screen include G protein-coupled receptors, ion channels, transporters, kinases, proteases, neurotransmitters, phosphodiesterases, and nuclear hormone receptors (total 87 safety screens)^32^. The methods used were reported to be radioligand-based binding and enzymatic biochemical assays. The study was executed by Eurofins (France). The list of assays, associating ligands/ stimulus and concentration are shown in Supplementary Table 5 and Supplementary Table 6. SP16 was tested at 10 µM. The results are expressed as a percent inhibition of control specific binding.

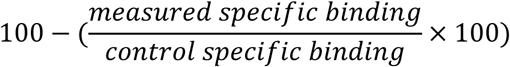

Results showing an inhibition or stimulation higher than 50% are considered to represent significant effects of the test compounds.

### Surface plasmon resonance

Biotinylated SP16 and SP34 (SP16 scrambled) were immobilized onto streptavidin coated Sensor Chip SA (Cytiva, BR100531) using running buffer containing HEPES (pH 7.4), 150 mM NaCl (Merck, S5886) and 0.05% Tween20 (Sigma, P9416-50ML) at flow rate of 10 µL/min and concentration of 1 µM peptides. Recombinant proteins of human insulin receptor/ CD220 (28-944) protein, His Tag (ACROBiosystems, INR-H52Ha) were injected over the peptide coupled sensor (concentration from 0.7 nM to 500 nM). Multicycle kinetics injections were performed in running buffer at a flow rate of 30 μL/min with an association and dissociation time of 210 s and 600 s, respectively, and binding was measured in relative response units. Data analysis was performed byusing Biacore T200 Evaluation Software 3.2. Sensorgrams were fitted with a 1:1 binding model using the Biacore Evaluation software and exported to GraphPad Prism 10.1.2.

### Competitive radioligand binding assay

A radioligand displacement assay was conducted using ChemiSCREEN Membrane Preparation Recombinant Human GIP Glucagon Family Receptor (Millipore, GIPR HTS112M SC845007) and Radioligand ^125^I GIP (Biotrend, A3-AI-745-T-100) following the manufacturer’s protocol. On the day prior, an FC 96-well harvest plate (Millipore) was coated with 0.33% polyethyleneimine and left overnight. The following day, the coated plate was washed once with wash buffer (50 mM HEPES pH 7.4, 500 mM NaCl). A mixture of 5 µg membrane, 25 pM radioligand and compounds was prepared in binding buffer containing 50 mM HEPES, pH 7.4, 5 mM MgCl2 (Sigma, M8266-100G), 1 mM CaCl2 (Merck,1023921000), 0.1% Ovalbumin (Serva, 11932), 0.01% Tween20 (Sigma, S4521-100). The mixture was incubated at room temperature for 4 h with shaking. The binding reaction was subsequently transferred to the pre-coated filter plate and washed 2 times (300 µL each) with wash buffer. The plate was air-dried at room temperature for 60 min. Next, 50 µL Microscint-20 (Millipore, 10084002) was added to each well, and radioactivity was counted with TOP Count NXT (PerkinElmer). IC50 and Ki were calculated with the GraphPad Prism software 10.1.2.

### Bulk mRNA sequencing

For acute treatment, the 9-day old differentiated hMADS cells were treated with DMSO 0.1%, forskolin (10 µM), GIP (10 nM), SP16 (10 µM), SP16 NC (10 µM) for 4 h (37 °C, 5% CO2).

Cells were lysed, and RNA was isolated. For treatment with insulin, 9-day old differentiated hMADS cells were starved with starvation medium overnight (37 °C, 5% CO2). The following day, hMADS cells were incubated with DMSO 0.1%, picropodophyllin 10 µM (PPP, Sigma, 407247), hGIP 10 nM, SP16 10 µM, SP16 NC 10 µM for 30 min. Insulin was added to the corresponding samples and incubated for another 24 h (37 °C, 5% CO2). After the incubation time, the cells were washed with PBS, lysed and RNA was isolated.

For bulk RNA-sequencing, total RNA derived from hMADS (n=6) was quantitatively and qualitatively assessed using the fluorescence Broad Range Quant-iT RNA Assay Kit (Thermo Fisher Scientific) and the Standard Sensitivity RNA Analysis DNF-471 Kit on a 96-channel Fragment Analyzer (Agilent), respectively. All total RNA samples had a RIN > 8. Total RNA input of 250 ng was employed for library preparation with the NEBNext UltraExpress RNA Library Prep Kit for Illumina (NEB, #E3330), together with the NEBNext Poly(A) mRNA Magnetic Isolation Module (NEB, #E7490) and NEBNext Multiplex Oligos for Illumina (NEB, #E7600) as per manufacturer’s instructions on a Biomek i7 (Beckman Coulter). For double-stranded cDNA purification, Ampure XP beads (Beckman Coulter) were used instead of the recommended SPRIselect Beads. Libraries were amplified with 13 PCR cycles. The final RNA-seq libraries were eluted in 30 µL EB Buffer (Qiagen) and were quantified by the High Sensitivity dsDNA Quanti-iT Assay Kit (ThermoFisher) on a Synergy HTX (BioTek). Libraries were assessed for size distribution and adapter dimer presence (<0.5%) by the High Sensitivity Small Fragment DNF-477 Kit on a 96-channel Fragment Analyzer (Agilent). Libraries were normalized on the MicroLab STAR (Hamilton), pooled and sequenced on a NovaSeq 6000 (Illumina) with dual index, paired-end reads at 2 x 100 bp length (Rd1: 101bp, Rd2: 8bp, Rd3: 8bp, Rd4:101bp) with an average sequencing depth of ∼32 million Pass-Filter reads per sample.

Sequencing reads from the RNA-seq experiment were processed with a pipeline built upon the implementation of the ENCODE’ “Long RNA-seq” pipeline: Filtered reads were mapped against the Homo sapiens (humans) genome hg38/GRCh38 (primary assembly, excluding alternate contigs) using the STAR aligner software^54^ (STAR version 2.5.2b) allowing for soft clipping of adapter sequences. For quantification of transcript levels, the annotation files from Ensembl version 86 were used, which corresponds to GENCODE 25. Samples were quantified with the above annotations, using RSEM^55^ (RSEM version 1.3.0) and featureCount^56^ (featureCount version 1.5.1). Quality controls were implemented using FastQC^57^ (FastQC version 0.11.5), picardmetrics (picardmetrics version 0.2.4), and dupRadar^58^ (dupRadar version 1.0.0) at the respective steps. Differential expression analysis was performed on the mapped counts derived from featureCount using limma/voom^59,60^. Furthermore, false discovery rate (FDR) of ≤ 0.05 was used for further analysis. To derive the pathways containing differentially regulated genes. We used the clusterProfiler^9^ R package on the database Gene-Ontology (GO), ReactomePA (1.36.0)^61^ and gene set enrichment analysis was performed, using log Fold-Change as ranking metric^11^.

### Proteomics

#### Sample preparation

Sample preparation was performed in a 96-well format (twin.tec PCR Plate 96 LoBind, Eppendorf) following an adjusted in-StageTip protocol^62^. For plasma sample preparation, 20 µL of PreOmics Enrich beads (PreOmics Enrich Kit) were added to a 96-deep well round plate (Eppendorf, 0030504100) washed utilizing 100 µL of beads wash buffer (PreOmics Enrich Kit), followed by incubation at 30 °C for 1 min on a thermo shaker (ThermoMixer C, Eppendorf) with a subsequent bead precipitation on a magnet plate (Alpaqua Magnum FLX 96-well microplate) for 1 min. In total, beads were washed three times. Subsequently, 20 µL of plasma and 40 µL of binding buffer (PreOmics Enrich Kit) were added to the beads, followed by a 30 min incubation at 1,200 rpm, 30 °C to enable low abundant proteins to bind to the bead slurry. Then, the supernatant was removed and 100 µL of binding buffer was added to repeat the wash step two more times.

To continue with either tissue or plasma preparation the PreOmics iST 192x kit was used (PreOmics, catalogue number P.O.00067) and samples were processed according to manufacturer’s instructions. First, 50 µL of lysis buffer were added to the washed beads binding plasma proteins and samples were incubated for 20 min at 80 °C and 1,200 rpm on a thermo shaker. Samples were cooled down to room temperature and 50 µL of Trypsin/LysC mixture was added to the sample, followed by overnight incubation at 37 °C and 1,200 rpm on a thermo shaker. Next day, sample digestion was stopped by adding 100 µL of stop buffer (PreOmics iST kit), followed by a transfer into the provided solid phase extraction (SPE) 96- well plate. The SPE plate was positioned on top of an Eppendorf 96-deep well round plate and centrifuged for 2 min at 2,200 g on a benchtop centrifuge. Flowthrough was discarded, 200 µL of Wash 1 buffer (PreOmics iST kit) were added and centrifugation repeated. This wash step was repeated one more time. Then, 200 µL of Wash 2 buffer (PreOmics iST kit) were added and centrifugation repeated. This step was repeated one more time. The SPE containing the clean peptides was positioned on an Eppendorf 96-well PCR plate, 40 µL of provided elution buffer (Elute, PreOmics iST kit) were added and centrifugation repeated. This step was repeated to collect a total of 80 µL flowthrough containing the cleaned-up peptides. Eluted, clean peptides were dried (Concentrator plus, Eppendorf) and resuspended in 25 µL of LC-load buffer (PreOmics iST kit) with a 30 min incubation step at 30°C, 1200 rpm. The peptide concentration per well was quantified utilizing a colorimetric quantitative peptide assay (catalogue number 23275, Pierce, Quantitative Peptide Assay, Thermo Fisher Scientific).

#### LC-MS analysis

Proteome analysis was performed utilizing the Vanquish Neo system (Thermo Fisher Scientific) coupled to an Orbitrap Astral mass spectrometer (Thermo Fisher Scientific) with a nano-electrospray source. 500 ng of each sample was injected and separated with a 15 cm IonOpticks Aurora Elite XT C18 column (75 µm inner diameter, 150 mm length, IonOpticks) heated to 50°C and run at a 500 nL/min flow rate. The chromatography gradient was set to 30 min total, separating peptides by hydrophobicity for 25 min starting with 1% Buffer B (80% ACN (Acetonitrile) 19.9% ddH2O (double-distilled water), 0.1% FA (Formic Acid)), 99% Buffer A (99.9% ddH2O, 0.1% FA) and ending with 30% Buffer B and 70% Buffer A. For column wash and regeneration, Buffer B was ramped up to 95% in 1 min and kept for 4 min at 95% Buffer B until the end of the gradient. Column regeneration was performed automatically before sample pickup for the next run. Electrospray voltage was set to 1750 V in positive ion mode. Mass spectrometry analysis was performed using a data independent acquisition (DIA) scan mode.

#### Upstream data processing

Mass spectrometry raw data were processed within the DIA-NN software suite^63^ (utilizing the directDIA search algorithm. Searches were performed utilizing standard settings except for: For precursor ion generation “Fasta digest for library-free search/library generation” and “Deep learning-based spectra, retention times, and ion mobility value prediction” was checked; Precursor m/z range was set to cover 350-1400; Fragment ion m/z range was set to cover 150-2000; “Neural network classifier” was set to “Double-pass mode”. Searches were performed against the mus musculus UniProt reference proteome of canonical and isoform sequences (UP00000589_10090.fa, UP00000589_10090_additional.fa).

#### Downstream data processing

Plasma proteomics data were filtered for at least 4,000 protein identifications and a coefficient of variation (CV) of equal or less than two and at least 70% data completeness in at least one group for improved data consistency. Liver proteomics data were not filtered for any additional data consistency criteria. Missing values were imputed from a down-shifted normal distribution by 1.8 and a width of 0.3 standard deviations for both biospecimen. Differential expression analysis was performed using a two-sided t-test utilizing a false-discovery rate of 5% and 250 randomized tests to correct for multiple hypothesis testing including a S0-modulation of 0.1. Correlation analysis was performed utilizing Pearson correlation as a metric.

#### *In vivo* experiments

The experimental protocols involving the use of laboratory animals were reviewed and approved by the relevant governmental authorities. The mice were housed under 12 h light - dark cycle and had *ad libitum* access to food (Haltungsfutter Altromin1328 for PK study and normal chow diet, and HFD D12492, Research Diets, Inc., New Brunswick, NJ, USA for DIO mice study) and tap water, unless stated otherwise. The vehicle employed for the duration of the experiments consisted of a 20 mM acetate buffer pH 4, supplemented with 5% mannitol (Merck, M0200000).

### Pharmacokinetic properties of SP16

C57BL/6J male mice purchased from The Jackson Laboratories were used. A single intraperitoneal (i.p.) administration of SP16 was given to each mouse (n=3) at a dose of 300 nmol/kg, with an application volume of 10 mL/kg. Blood samples were obtained at baseline, 5, 15, 30, 60, 120, and 240 min, and at 8 h and 24 h after administration. Plasma samples were generated from these blood samples and stored at -20 °C until further analysis. To measure the plasma concentrations of SP16, a liquid chromatography/tandem mass spectrometry (LC-MS/MS) method was employed using a QTRAP 6500+ instrument (Sciex, Framingham, MA, USA). The calibration range of the method was set from 0.5 to 1000 nM. Prior to analysis, the plasma samples were subjected to protein precipitation by treating them with ethanol.

### Oral glucose tolerance test

To assess the ability of SP16 to modulate glucose tolerance, an oral glucose tolerance test (oGTT) was performed. C57BL/6J male mice fed with a 60% high fat diet (HFD) were purchased from The Jackson Laboratories at >16 weeks of age. The administration of the glucose bolus was designated as 0 h. The mice were fasted at -12 h and randomized based on weight into 4 groups at -6 h (n = 8 mice per group). Mice were administered s.c. GIPR agonist (patent number: WO/2018/181864) at -4 h, or intraperitoneal (i.p.) SP16 at 0 h. Baseline blood glucose was measured at -30 min. Oral bolus of 2 g/kg glucose was applied at 0 h. Blood glucose was measured at 0, 15, 30, 60, and 120 min using a glucometer (GlucoSmart Swing -10032).

### Acute treatment of SP16 on DIO mice and lipolysis on tissue

After the oGTT experiment, the mice were allowed to recover for at least one week while continuing the HFD. On the day of experiment, mice treated with vehicle or SP16 (application volume 5 mL/kg) were sacrificed 15 min after i.p. administration, while mice treated with GIPR agonist (patent number: WO/2018/181864) were sacrificed 4 h after subcutaneous (s.c). administration. The mice were sedated with isoflurane. Blood was drawn from retro-orbital sinus and treated with EDTA as an anticoagulant. iWAT, eWAT, iBAT, and liver were collected. Each tissue was cut into small pieces weighing of ca. 10 - 20 mg and placed into pre-warmed KRBB supplemented with 2% BSA for the lipolysis assay. The remaining harvested tissues were snap-frozen in liquid nitrogen and stored at -80 °C for further analysis. Plasma triglycerides (TG), total cholesterol (TC), aspartate aminotransferase (AST), and alanine aminotransferase (ALT) were determined using the Cobas measurement system. FFA (Wako), glycerol (Sigma, F6428-40ML), leptin (Millipore, RAB0334-1KT), IL-1β (Millipore, RAB0273), TNFα (Millipore, RAB0476-1KT) were measured following the manufacturer’s protocol. cAMP concentration in tissue was measured with cAMP ELISA as previously described. Phosphorylated AKT levels were determined using Simple Western Blot (ProteinSimple).

### Data and statistical analyses

The data analysis was performed using GraphPad Prism 10 and Microsoft Excel. The data is presented as mean ± standard error of the mean (SEM) from individual measurements. To compare two groups with one variable, one-way ANOVA multiple comparison was utilized. For experiments comparing two groups and two variables, a two-way ANOVA multiple comparison was conducted. Unless stated otherwise, p-values less than 0.05 between two groups were deemed significant and indicated with asterisk(s), while non-significant comparisons had no indication.

## Acknowledgement

We thank Anke Voigt and Juliane Denz for performing *in vivo* experiment. We thank Yvette Hoevels for performing SPR assay. We thank Sandra Gross for performing radioligand binding assay, Werner Rust for performing RNA sequencing, and Frank Wesche for PK analysis. We thank Victoria Schröder, Tobias Kiechle, and Kimberly Thomas for giving help in cell culturing. We thank Bradford Hamilton, Sandra Kleiner, Daniel Markgraf, and Tim Kloekener for their input. We thank Marco Block and Alvaro Carcamo Martinez for helping with extended-release formulation. This work was supported by Boehringer Ingelheim Pharma & Co. KG.

## Contribution

Thi My Hanh Ngo: Conceptualization, methodology, investigation, formal analysis, data curation, validation, visualization, writing – original draft.

Rakesh Santhanam.: RNA data analysis, formal analysis, data curation, visualization

Daniel P. Teufel: Peptide synthesis, methodology, conceptualization, writing – review and editing

Alec Dick: RNA sequencing, writing – review and editing Daniel Lam: RNA data analysis, writing – review and editing Holger Klein: RNA data analysis, writing – review and editing

Andreas-David Brunner: Proteome analysis, Visualization, writing – review and editing

Mafalda M.A. Pereira: *in vivo* study, methodology, data curation, validation, visualization, writing – review and editing

Alexander Bartelt: Conceptualization, writing – review and editing Anton Pekcec: Conceptualization, writing – review and editing

Maude Giroud: Conceptualization, supervision, project administration, writing – review and editing

## References

1. Fruh, S. M. Obesity: Risk factors, complications, and strategies for sustainable long-term weight management. J. Am. Assoc. Nurse Pr. 29, S3–S14 (2017).

2. Sarwer, D. B. & Polonsky, H. M. The Psychosocial Burden of Obesity. Endocrinol. Metab. Clin. North Am. 45, 677–688 (2016).

3. Yanovski, S. Z. & Yanovski, J. A. Long-term Drug Treatment for Obesity: A Systematic and Clinical Review. Jama 311, 74–86 (2014).

4. Holst, J. J. The incretin system in healthy humans: The role of GIP and GLP-1. Metabolism 96, 46–55 (2019).

5. Holst, J. J. & Rosenkilde, M. M. GIP as a Therapeutic Target in Diabetes and Obesity: Insight From Incretin Co-agonists. J. Clin. Endocrinol. Metab. 105, dgaa327 (2020).

6. Killion, E. A. et al. Glucose-Dependent Insulinotropic Polypeptide Receptor Therapies for the Treatment of Obesity, Do Agonists = Antagonists? Endocr. Rev. 41, bnz002 (2019).

7. Regmi, A. et al. Tirzepatide modulates the regulation of adipocyte nutrient metabolism through long-acting activation of the GIP receptor. Cell Metab. 36, 1534–1549.e7 (2024).

8. Xiao, J., Kim, S.-J., Cohen, P. & Yen, K. Humanin: Functional Interfaces with IGF-I. 29, 21– 27 (2016).

9. Boutari, C., Pappas, P. D., Theodoridis, T. D. & Vavilis, D. Humanin and diabetes mellitus: A review of in vitro and in vivo studies. World J. Diabetes 13, 213–223 (2022).

10. Cui, X. et al. The secreted peptide BATSP1 promotes thermogenesis in adipocytes. Cell. Mol. Life Sci. 80, 377 (2023).

11. Yu, X. et al. The GIP receptor activates futile calcium cycling in white adipose tissue to increase energy expenditure and drive weight loss in mice. Cell Metab. (2024) doi:10.1016/j.cmet.2024.11.003.

12. Serres, F. de & Blanco, I. Role of alpha-1 antitrypsin in human health and disease. J. Intern. Med. 276, 311–335 (2014).

13. Stocks, B. et al. Integrated Liver and Plasma Proteomics in Obese Mice Reveals Complex Metabolic Regulation. Mol. Cell. Proteom. 21, 100207 (2022).

14. Mansuy-Aubert, V. et al. Imbalance between Neutrophil Elastase and its Inhibitor α1- Antitrypsin in Obesity Alters Insulin Sensitivity, Inflammation, and Energy Expenditure. Cell Metab. 17, 534–548 (2013).

15. Bigalke, A., Sponholz, C., Schnabel, C., Bauer, M. & Kiehntopf, M. Multiplex quantification of C-terminal alpha-1-antitrypsin peptides provides a novel approach for characterizing systemic inflammation. Sci. Rep. 12, 3844 (2022).

16. Lior, Y., Shtriker, E., Kahremany, S., Lewis, E. C. & Gruzman, A. Development of anti-inflammatory peptidomimetics based on the structure of human alpha1-antitrypsin. Eur. J. Med. Chem. 228, 113969 (2022).

17. Wang, Z. et al. α1-Antitrypsin derived SP16 peptide demonstrates efficacy in rodent models of acute and neuropathic pain. Faseb J 36, e22093 (2022).

18. Toldo, S. et al. Low-Density Lipoprotein Receptor–Related Protein-1 Is a Therapeutic Target in Acute Myocardial Infarction. Jacc Basic Transl Sci 2, 561–574 (2017).

19. Madsen, C. T. et al. Combining mass spectrometry and machine learning to discover bioactive peptides. Nat. Commun. 13, 6235 (2022).

20. Arapidi, G. et al. Peptidomics dataset: Blood plasma and serum samples of healthy donors fractionated on a set of chromatography sorbents. Data Brief 18, 1204–1211 (2018).

21. Zhang, X., Ostrov, D. A. & Tian, H. Alpha-1 antitrypsin: A novel biomarker and potential therapeutic approach for metabolic diseases. Clin. Chim. Acta 534, 71–76 (2022).

22. Wohlford, G. F. et al. A phase 1 clinical trial of SP16, a first-in-class anti-inflammatory LRP1 agonist, in healthy volunteers. Plos One 16, e0247357 (2021).

23. Liu, Q. et al. Lipoprotein Receptor LRP1 Regulates Leptin Signaling and Energy Homeostasis in the Adult Central Nervous System. Plos Biol 9, e1000575 (2011).

24. Okagawa, S. et al. Hepatic SerpinA1 improves energy and glucose metabolism through regulation of preadipocyte proliferation and UCP1 expression. Nat. Commun. 15, 9585 (2024).

25. Shelbayeh, O. A., Arroum, T., Morris, S. & Busch, K. B. PGC-1α Is a Master Regulator of Mitochondrial Lifecycle and ROS Stress Response. Antioxidants 12, 1075 (2023).

26. Markussen, L. K. et al. Lipolysis regulates major transcriptional programs in brown adipocytes. Nat. Commun. 13, 3956 (2022).

27. Bartelt, A. & Heeren, J. Adipose tissue browning and metabolic health. Nat. Rev. Endocrinol. 10, 24–36 (2014).

28. Cero, C., et al. β3-Adrenergic receptors regulate human brown/beige adipocyte lipolysis and thermogenesis. JCI Insight 6, e139160 (2021).

29. Müller, T. D. et al. Glucose-dependent insulinotropic polypeptide (GIP). Mol. Metab. 95, 102118 (2025).

30. Yang, B. et al. Discovery of a potent GIPR peptide antagonist that is effective in rodent and human systems. Mol. Metab. 66, 101638 (2022).

31. Tallarida, R. J. & Murray, R. B. pA2 Analysis I: Schild Plot. in (eds. Tallarida", ["Ronald J. & Murray"], "Rodney B.) 53–56 (Springer New York, New York, NY, 1987). doi:10.1007/978-1-4612-4974-0_16.

32. Brennan, R. J. et al. The state of the art in secondary pharmacology and its impact on the safety of new medicines. Nat. Rev. Drug Discov. 23, 525–545 (2024).

33. Lee, J. & Pilch, P. F. The insulin receptor: structure, function, and signaling. American Journal of Physiology-Cell Physiology 266, C319–C334 (1994).

34. Kahn, C. R. & White, M. F. The insulin receptor and the molecular mechanism of insulin action. J. Clin. Investig. 82, 1151–1156 (1988).

35. Cignarelli, A. et al. Insulin and Insulin Receptors in Adipose Tissue Development. Int. J. Mol. Sci. 20, 759 (2019).

36. Campbell, J. E. Targeting the GIPR for obesity: To agonize or antagonize? Potential mechanisms. Mol. Metab. 46, 101139 (2021).

37. Zhang, Y. et al. Comparative Proteomic Analysis of Liver Tissues and Serum in db/db Mice. Int. J. Mol. Sci. 23, 9687 (2022).

38. Coleman, B. A. & Taylor, P. Regulation of Acetylcholinesterase Expression during Neuronal Differentiation (∗). J. Biol. Chem. 271, 4410–4416 (1996).

39. Walczak-Nowicka, Ł. J. & Herbet, M. Acetylcholinesterase Inhibitors in the Treatment of Neurodegenerative Diseases and the Role of Acetylcholinesterase in their Pathogenesis. Int. J. Mol. Sci. 22, 9290 (2021).

40. Soler, Y. et al. SERPIN-Derived Small Peptide (SP16) as a Potential Therapeutic Agent against HIV-Induced Inflammatory Molecules and Viral Replication in Cells of the Central Nervous System. Cells 12, 632 (2023).

41. Ding, Y., Xian, X., Holland, W. L., Tsai, S. & Herz, J. Low-Density Lipoprotein Receptor-Related Protein-1 Protects Against Hepatic Insulin Resistance and Hepatic Steatosis. Ebiomedicine 7, 135–145 (2016).

42. Kang, M.-C. et al. LRP1 regulates food intake and energy balance in GABAergic neurons independently of leptin action. Am J Physiol-endoc M 320, E379–E389 (2021).

43. Chandrasekaran, P. & Weiskirchen, R. Cellular and Molecular Mechanisms of Insulin Resistance. Curr. Tissue Microenviron. Rep. 5, 79–90 (2024).

44. Wang, J., Dai, L., Yu, T. & Xiao, J. Insights into the progressive impact of high-fat-diet induced insulin resistance on skeletal muscle and myocardium: A comprehensive study on C57BL6 mice. PLOS ONE 20, e0310458 (2025).

45. Lang, P., Hasselwander, S., Li, H. & Xia, N. Effects of different diets used in diet-induced obesity models on insulin resistance and vascular dysfunction in C57BL/6 mice. Sci. Rep. 9, 19556 (2019).

46. Gasbjerg, L. S. et al. Altered desensitization and internalization patterns of rodent versus human glucose-dependent insulinotropic polypeptide (GIP) receptors. An important drug discovery challenge. Br. J. Pharmacol. (2024) doi:10.1111/bph.16478.

47. Ceperuelo-Mallafré, V. et al. Disruption of GIP/GIPR Axis in Human Adipose Tissue Is Linked to Obesity and Insulin Resistance. J. Clin. Endocrinol. Metab. 99, E908–E919 (2014).

48. Lynggaard, M. B., Gasbjerg, L. S., Christensen, M. B. & Knop, F. K. GIP(3-30)NH2 – a tool for the study of GIP physiology. Curr. Opin. Pharmacol. 55, 31–40 (2020).

49. Block, M. et al. Miniaturized screening and performance prediction of tailored subcutaneous extended-release formulations for preclinical in vivo studies. Eur. J. Pharm. Sci. 196, 106733 (2024).

50. Arapidi, G. et al. Peptidomics dataset: Blood plasma and serum samples of healthy donors fractionated on a set of chromatography sorbents. Data Brief 18, 1204–1211 (2018).

51. Giroud, M. et al. HAND2 is a novel obesity-linked adipogenic transcription factor regulated by glucocorticoid signalling. Diabetologia 64, 1850–1865 (2021).

52. Estrada-Gutierrez, G. et al. Isolation of Viable Adipocytes and Stromal Vascular Fraction from Human Visceral Adipose Tissue Suitable for RNA Analysis and Macrophage Phenotyping. JoVE e61884 (2020) doi:10.3791/61884.

53. Cheng, Z. et al. Luciferase Reporter Assay System for Deciphering GPCR Pathways. Curr. Chem. Genom. 4, 84–91 (2010).

54. Dobin, A. et al. STAR: ultrafast universal RNA-seq aligner. Bioinformatics 29, 15–21 (2013).

55. Li, B. & Dewey, C. N. RSEM: accurate transcript quantification from RNA-Seq data with or without a reference genome. Bmc Bioinformatics 12, 323–323 (2011).

56. Liao, Y., Smyth, G. K. & Shi, W. featureCounts: an efficient general purpose program for assigning sequence reads to genomic features. Bioinformatics 30, 923–930 (2014).

57. Andrews & Simon. Babraham bioinformatics-FastQC a quality control tool for high throughput sequence data. (2010).

58. Sayols, S., Scherzinger, D. & Klein, H. dupRadar: a Bioconductor package for the assessment of PCR artifacts in RNA-Seq data. Bmc Bioinformatics 17, 428 (2016).

59. Ritchie, M. E. et al. limma powers differential expression analyses for RNA-sequencing and microarray studies. Nucleic Acids Res 43, e47–e47 (2015).

60. Law, C. W., Chen, Y., Shi, W. & Smyth, G. K. voom: precision weights unlock linear model analysis tools for RNA-seq read counts. Genome Biol 15, R29–R29 (2014).

61. Yu, G. & He, Q.-Y. ReactomePA: an R/Bioconductor package for reactome pathway analysis and visualization. Mol Biosyst 12, 477–479 (2015).

62. Kulak, N. A., Pichler, G., Paron, I., Nagaraj, N. & Mann, M. Minimal, encapsulated proteomic-sample processing applied to copy-number estimation in eukaryotic cells. Nat. Methods 11, 319–324 (2014).

63. Demichev, V., Messner, C. B., Vernardis, S. I., Lilley, K. S. & Ralser, M. DIA-NN: neural networks and interference correction enable deep proteome coverage in high throughput. Nat. Methods 17, 41–44 (2020).

