## Supplementary material for "The obesity-linked peptide SP16 regulates adipocytes through GIP and insulin receptor"

### Supplementary Figures

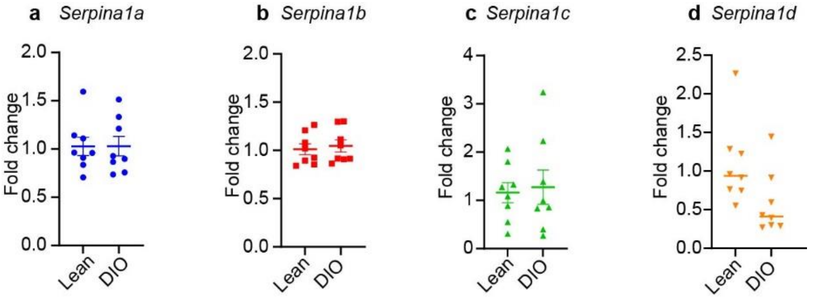


Supplementary Fig. 1: Expresson of *Serpina1* gene in lean and DIO mice. mRNA expression of *Serpina1a* (a), *Serpina1b* (b), *Serpina1c* (c), *Serpina1d* (d)

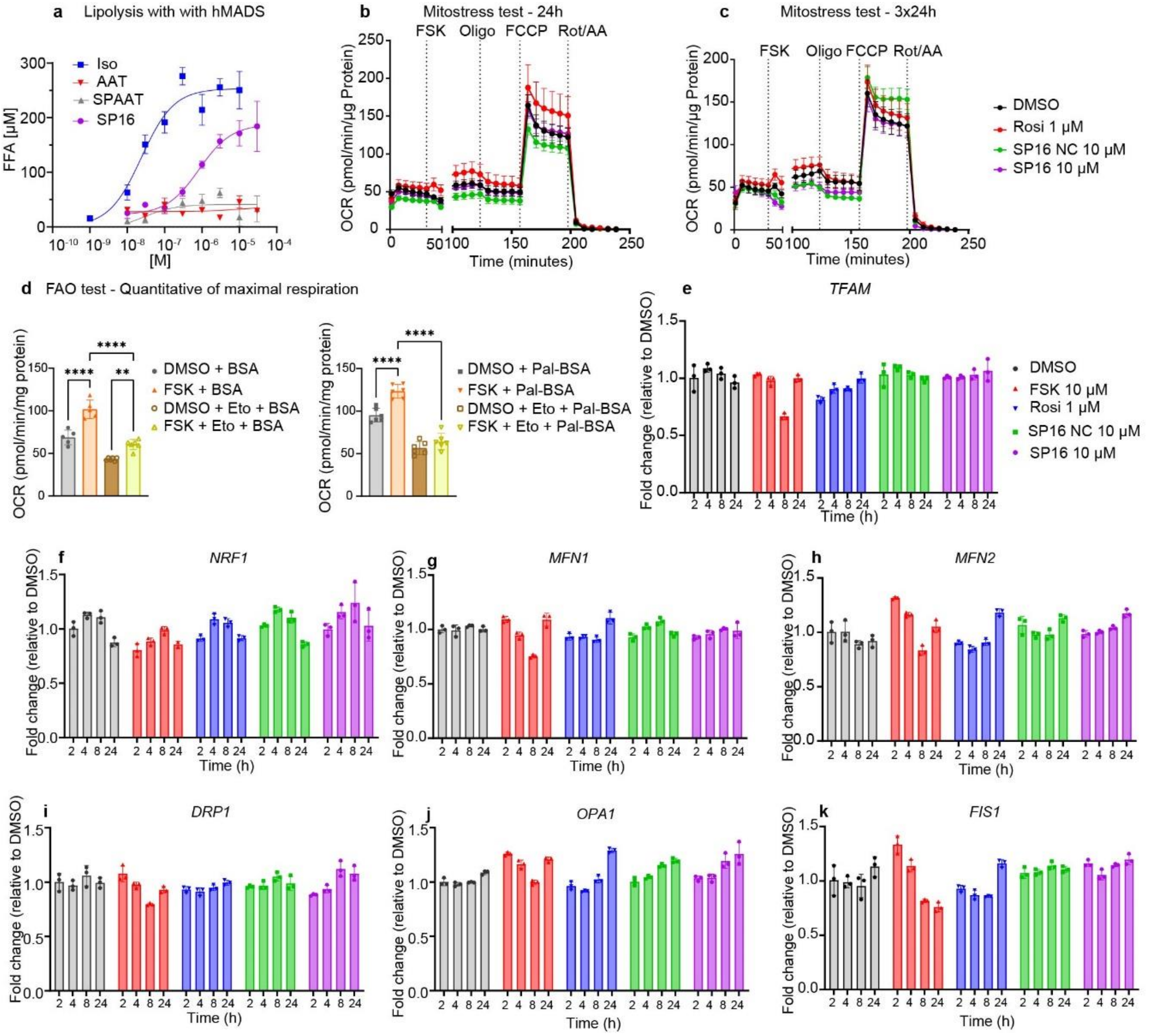


Supplementary Fig. 2. SP16 does not either induce OCR of hMADS when treated longer than 24 h or upregulate the expression of genes involving in biogenesis and dynamics of mitochondria. Lipolysis induced by AAT and peptides derived from AAT in hMADS (a), OCR of hMADS when treated with SP16 for 24 h (b) and 3x24 h (c), quantitative of maximal respiration of FAO stress test with DMSO and FSK as controls (d), mRNA expression of *TFAM* (e), *NFR1* (f), *MFN1* (g), *MFN2* (h), *DRP1* (i), *OPA1* (j), *FIS1* (k). Data are mean ± SEM, n=3-6, **p<0.01, ****p<0.0001 by one-way ANOVA with Dunnett's post-hoc test. OCR: oxygen consumption rate, FAO: fatty acid oxidation, Oligo: oligomycin, Rot/AA: rotenone/antimycin A, FSK: forskolin, SP16 NC: SP16 negative control, BSA: bovine serum albumin, Pal-BSA: palmitate conjugated with BSA, Eto: etomoxir, TFAM: mitochondrial transcription factor A, NRF1: nuclear respiratory factor 1, MFN1: mitofusin-1, MFN2: mitofusin-2, DRP1: dynamin-related protein 1, OPA1: optic atrophy 1, FIS1: fission-1.

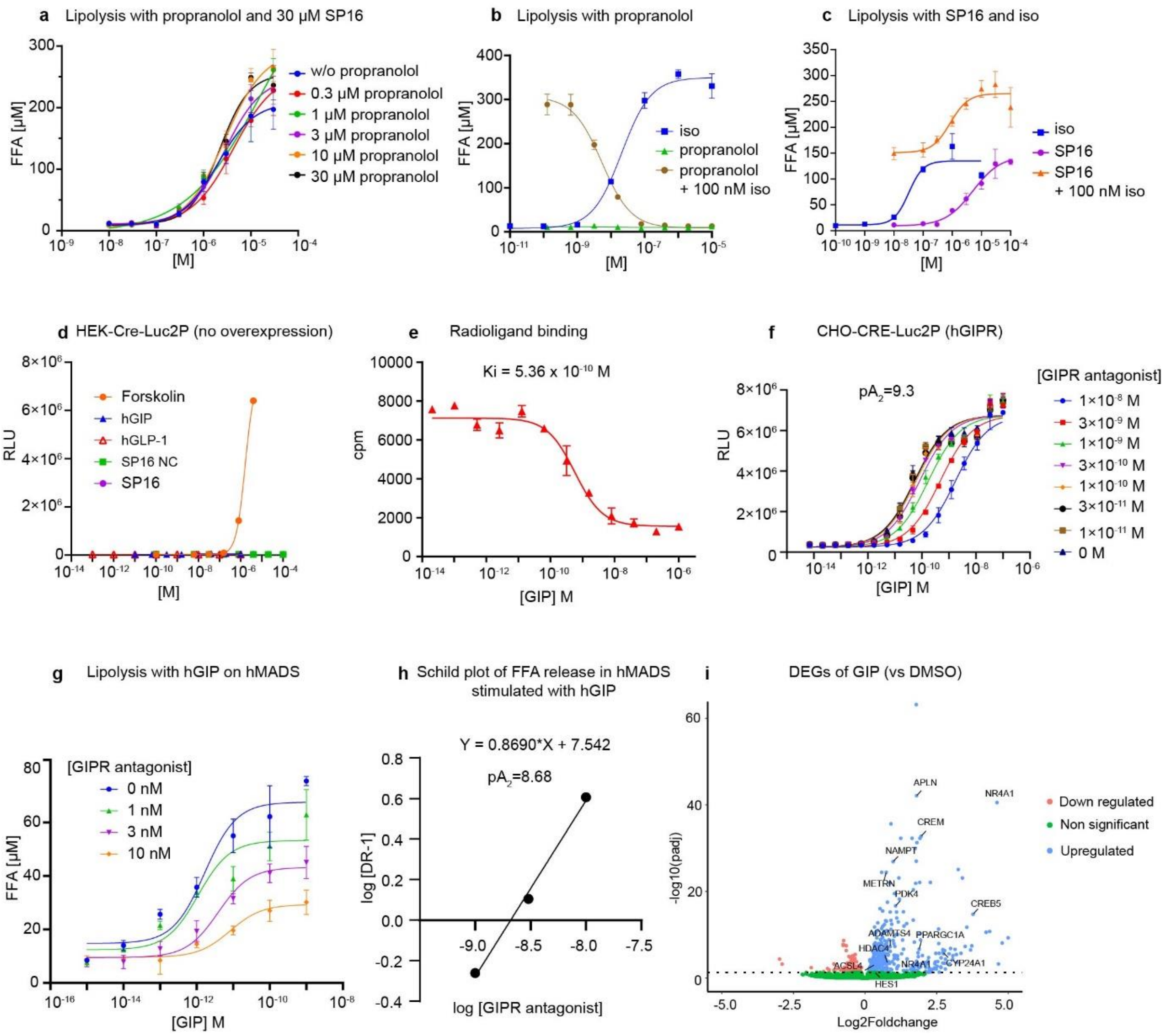


Supplementary Fig. 3: SP16 does not induce lipolysis through β-3 adrenergic receptora nd SP16 does not promote cAMP production in Cre-Luc2P reporter cells empty FFA release induced by SP16 when hMADS was pretreated with propranolol (a), FFA release induced by isoprenaline when hMADS was pretreated with propanolol (b), FFA release induced by SP16 or/and isoprenaline (c) luciferase activity in HEK-Cre-Luc2P report cells without receptor overexpression (d), competitive binding of hGIP to hGIPR membrane vs $^{125}$I-GIP (e), hGIP induced luciferase activity of CHO CRE-Luc2P cells overexpressing hGIPR pre-treated with different concentrations of GIPR antagonist (f), FFA release of hMADS induced by hGIP and different concentrations of GIPR antagonist (g), Schild plot obtained from FFA release of hMADS induced by hGIP when pretreated with GIPR antagonist (h), volcano plot of expression changes in GIP vs DMSO,abelled genes are involved in lipolysis, fatty acid activation, mitochondrial respiration and browning process (i) Data are mean ± SEM, n=4-8, RLU: relative light unit, FFA: free fatty acid.

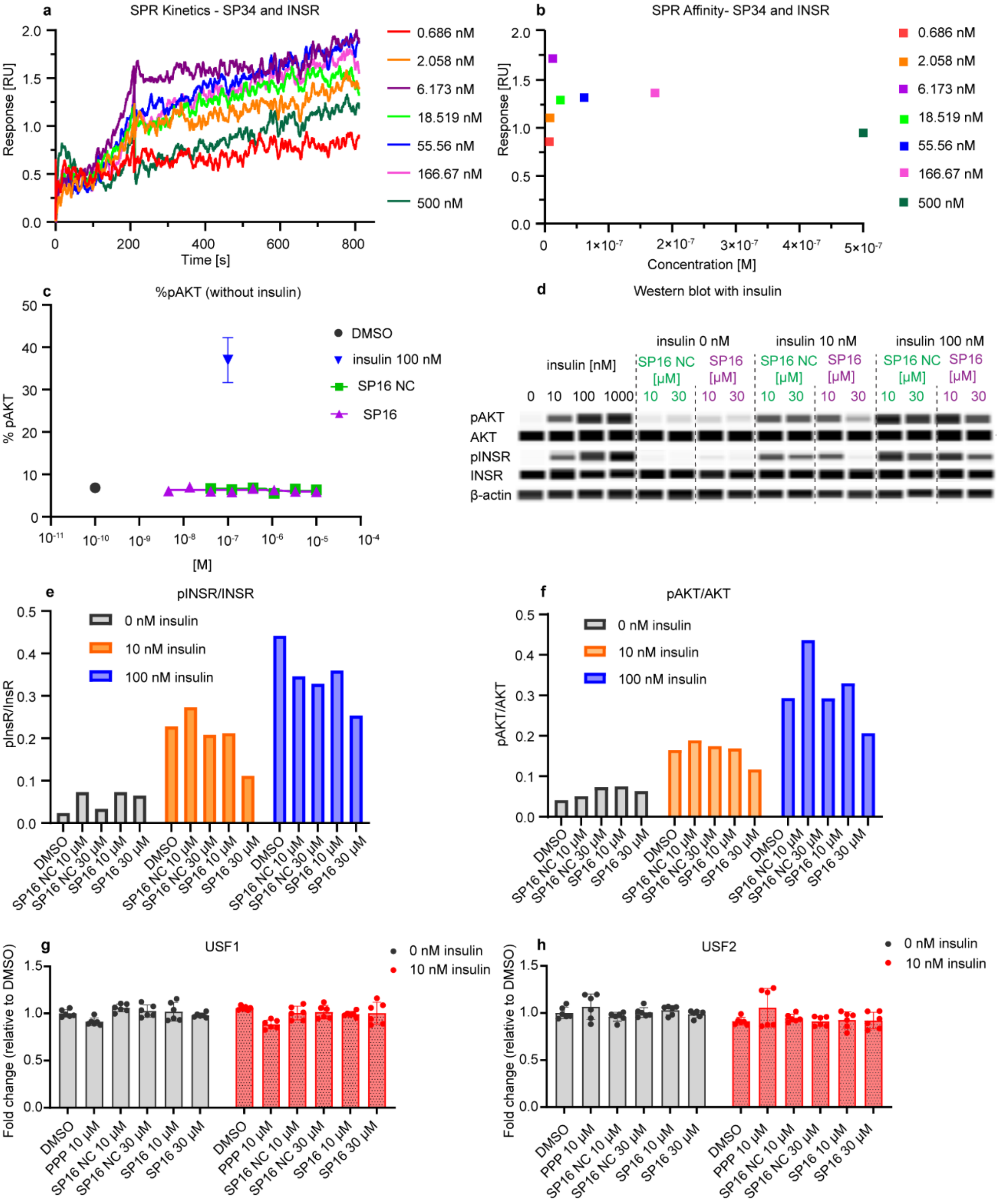


Supplementary Fig. 4: SP34 (SP16 scrambed) does not bind to INSR and SP16 does not modulate the expression of USF1 and USF2 in presence or absence of insulin. Kinetic fit of SPR binding assay between SP34 and INSR (a), affinity fit of SPR binding assay between SP34 and INSR (b), percentage of phosphorylated AKT of hMADS when treated with SP16 in absence of insulin (c), western blot of pAKT and pINSR of hMADS when treated with SP16 in absence of insulin and presence of insulin 10 nM and 100 nM (d), quantitative of pINSR/INSR (e), quantitative of pAKT/AKT (f), mRNA expression of USF1 (g) and USF2 (h) of hMADS when treated with insulin and SP16 for 24 h. Data are mean ± SEM, n=3-6. For (e) and (f), one data point is representative for the pool from 3 biological replicates. RU: resonance unit, INSR: insulin receptor, SP16 NC: SP16 negative control, PPP: picropodophyllin, USF1: upstream stimulatory factor 1, USF2: upstream stimulatory factor 2.

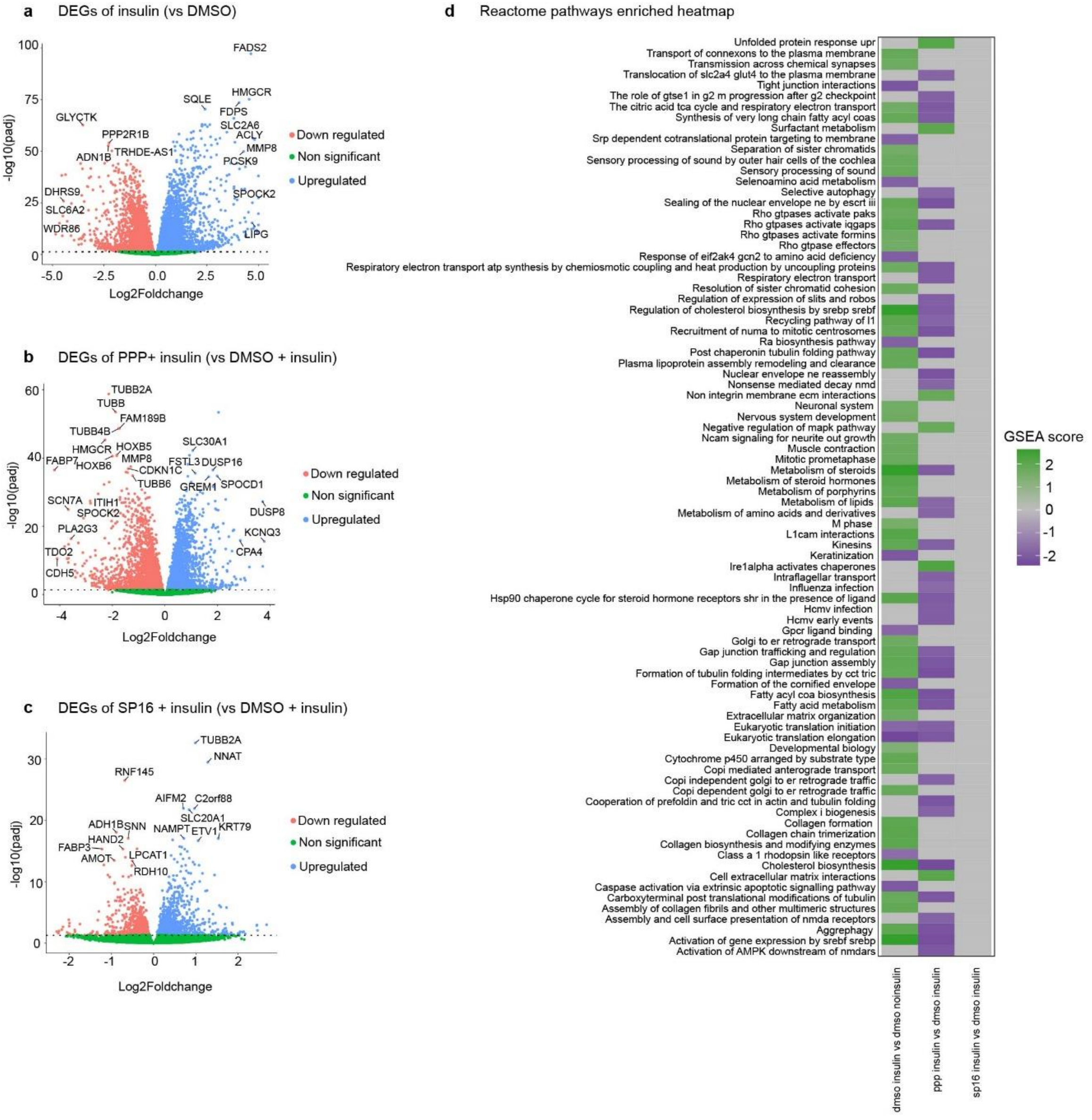


Supplementary Fig. 5: SP16 does not significantly interfere the metabolic pathways regulated by insulin in adipocytes. Volcano plots of DEGs (adj p < 0.05, FDR) of insulin (a), DEGs of PPP+ insulin (b), DEGs of SP16 + insulin (c), heatmap of Reactome pathways enriched by insulin treatment, PPP + insulin and SP16 + insulin, colur in tiles corresponds to GSEA score (d). DEG: differentially expressed genes, PPP: Picropodophylli.

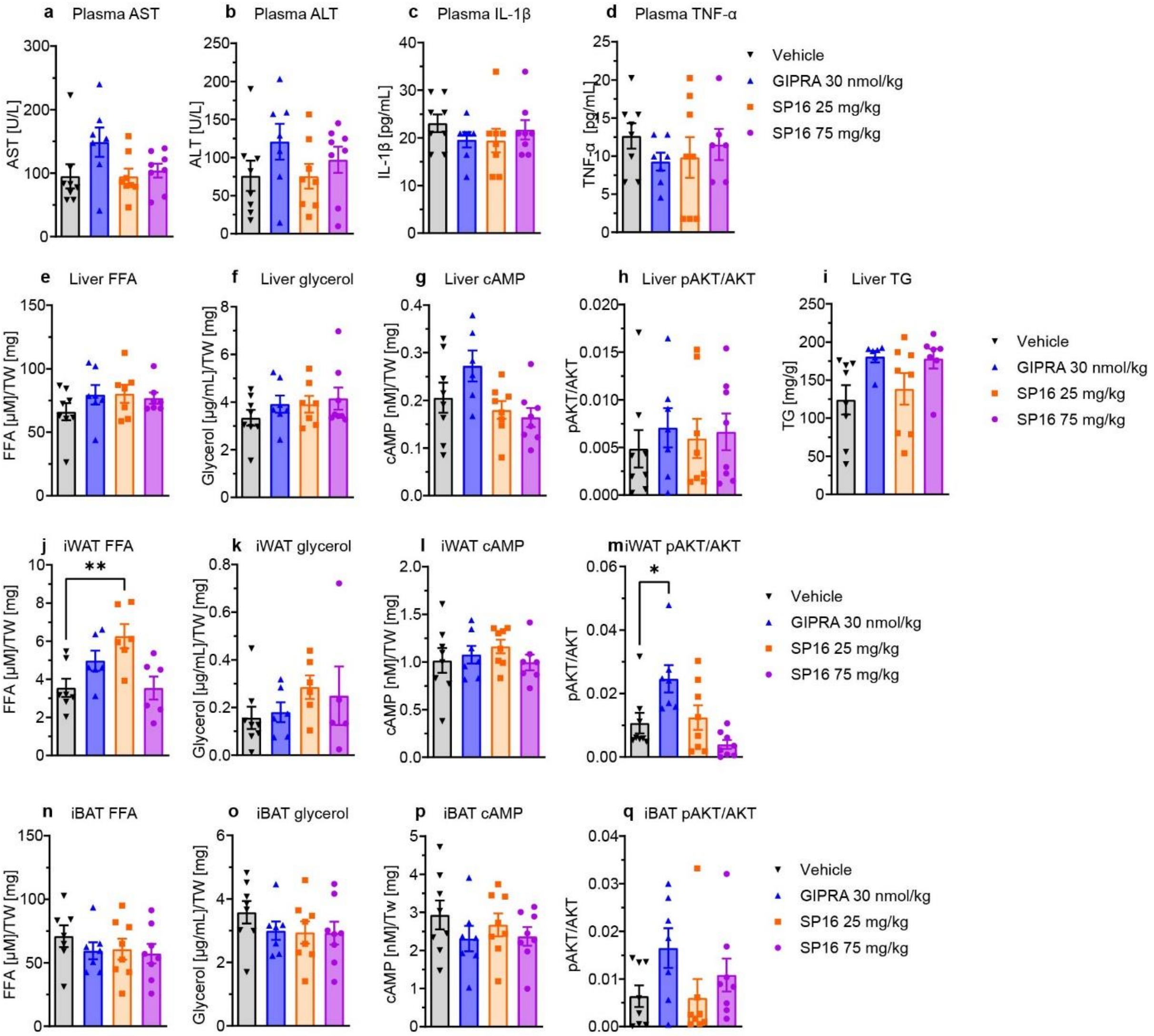


Supplementary Fig. 6: SP16 has short half-life, SP16 acute treatment in DIO mice slightly induce FFA release in iWAT and does not induce the FFA release, cAMP and pAKT in liver and BAT. Exposure of SP16 after subcutaneous injection of SP16 300 nmol/kg (a), plasma AST (b), plasma ALT (c), plasma IL-1β (d), plasma TNF-α (e), FFA release from liver (f), glycerol release from liver (g), cAMP level in liver (h), pAKT/AKT ratio in liver (i), total TG level in liver (j), FFA release from iWAT (k), glycerol release from iWAT (l), cAMP level in iWAT (m), pAKT/AKT ratio in iWAT (n), FFA release from iBAT (o), glycerol release from iBAT (p), cAMP level in iBAT (q), pAKT/AKT ratio in iBAT (r). Data are mean ± SEM, n=3-8, PK: pharmacokinetics, FFA: free fatty acid, TG: triglyceride, iWAT: inguinal white adipose tissue, iBAT: interscapular brown adipose tissue, TW: total weight of piece, GIPRA: GIPR agonist, AST: aspartat-aminotransferase, ALT: alanine aminotransferase.

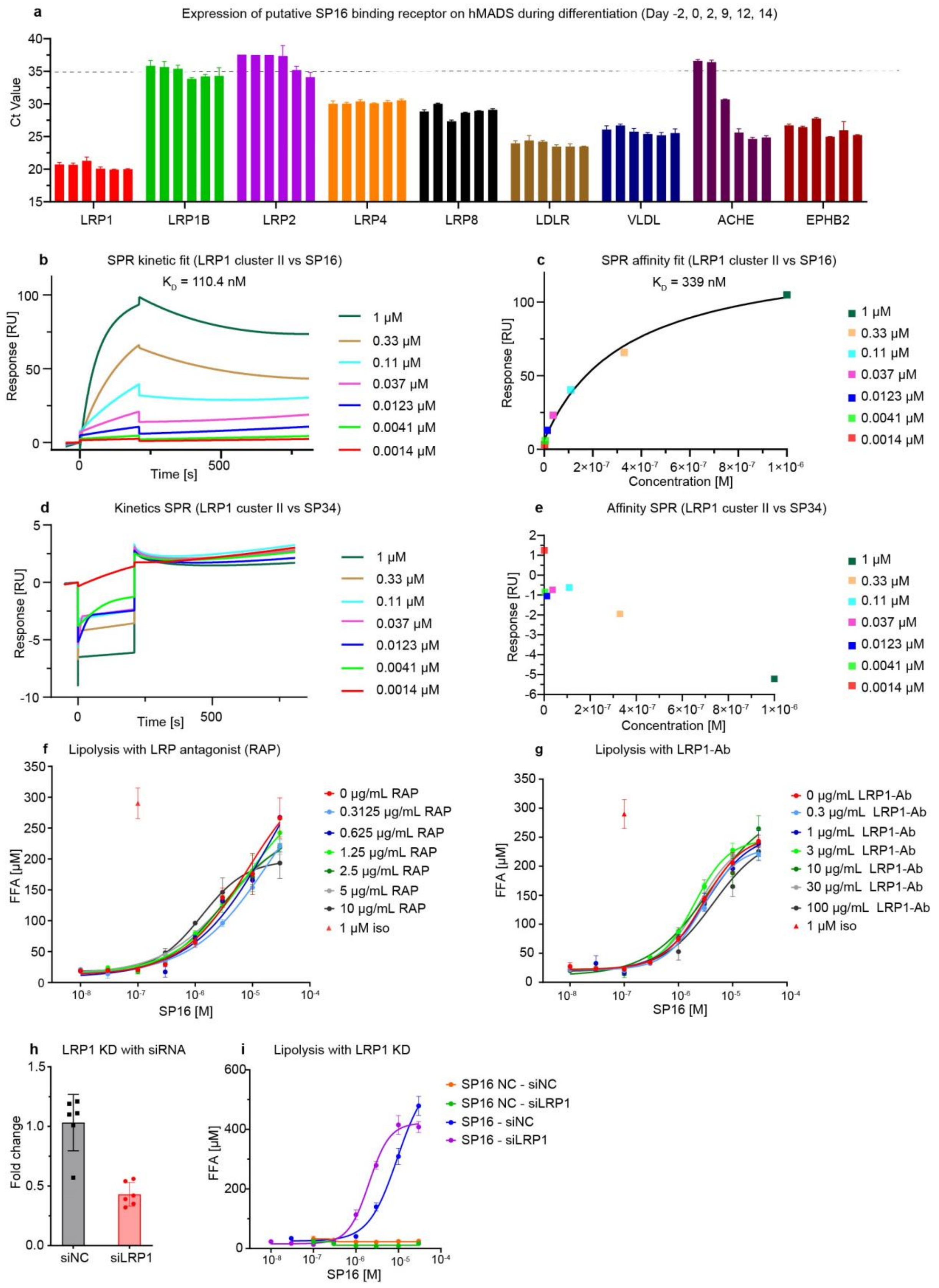


Supplementary Fig. 7: SP16 does not induce FFA of hMADS through LRP1 receptor. mRNA expression of putative SP16-binding receptors during differentiation , from left to right, day -2, day 0, day 2, day 9, day 14 (a), SPR kinetic fit of LRP1 to SP16 (b), SPR affinity fit of LRP1 to SP16 (c), SPR kinetic fit of LRP1 to SP34 (d), SPR affinity fit of LRP1 to SP34 (e), lipolysis of hMADS induced by SP16 and RAP (LRP antagonist) (f), or anti-LRP1-antibody (g), fold change in LRP1 gene expression in hMADS transfected with siRNA targeting GIPR or negative control siRNA (h), FFA release in cells transfected with siRNA targeting LRP1 or negative control siRNA (i), Data are mean ± SEM, n=3-8, RAP: receptor-associated protein, LRP1-Ab: anti-LRP1 antibody, KD: knockdown, siNC:siRNA negative control, siLRP1:siRNA targeting LRP1.

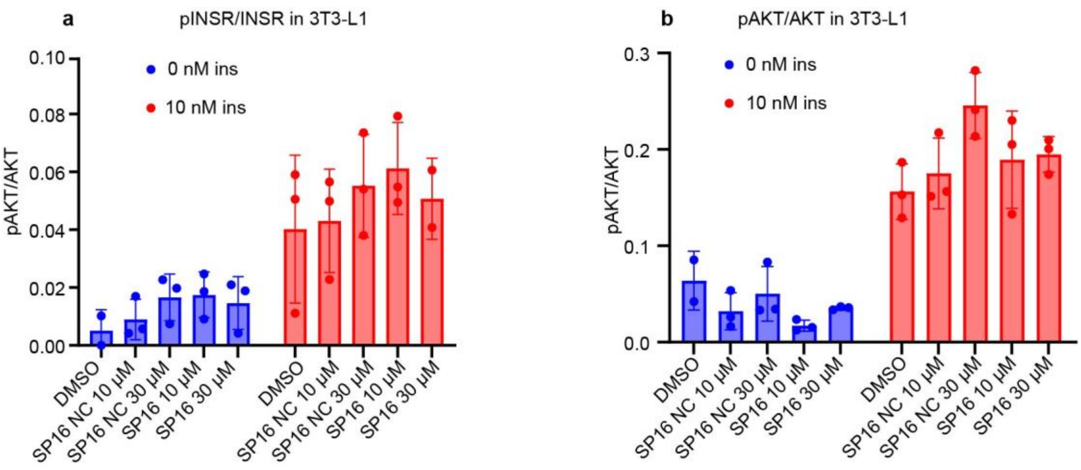


Supplementary Fig. 8: SP16 does not interferes the insulin signaling pathway in differentiated 3T3-L1 cells. pINSR/INSR induced by SP16 in absence and presence of 10 nM insulin (a), pAKT/AKT induced by SP16 in absence and presence of insulin 10 nM (b).

**Supplementary Tables**

Supplementary Table 1: Uniprot-annotated peptides derived from ProSAAS and AAT

| Protein | Sequence |
|---|---|
| ProSAAS | ARPVKEP |
| | ARPVKEPRGLSAASPPLAETGAPRRF |
| | GLSAASPPLAETGAPRRF |
| | AADHDVGSELPPEGVLGALLRV |
| | AADHDVGSELPPEGVLGALLRVKRLETPAPQVPARRLLPP |
| | LETPAPQVPA |
| | LETPAPQVPARRLLPP |
| AAT | MFLEAIPMSIPPEVKFNKPFVFLMIEQNTKSPLFMGKVVNPTQK |

Supplementary Table 2: Amino acid sequence of SPAAT, SP16, SP16 NC, SP34 and hGIP

| Peptide | Sequence |
|---|---|
| SPAAT | MFLEAIPMSIPPEVKFNKPFVFLMIEQNTKSPLFMGKVVNPTQK |
| SP16 | Ac-VKFNKPFVFL-Nle-IEQNTK-$NH_2$ |
| SP16 NC | Ac-VKFNKPAAAAAIEQNTK-$NH_2$ |
| SP34 | Ac-PKMVPQFNTELKIFPEVNIK-$NH_2$ |
| GIP (human GIP 1-42) | H-YAEGTFISDYSIAMDKIHQQDFVNWLLAQKGKKNDWKHNITQ-OH |

Peptides are N-terminally acetylated (“Ac-“) and C-terminally amidated (“-NH2). “Nle” refers to L-norleucine. All other amino acids are denoted by their one letter codes and are in their L-forms.

Supplementary Table 3**:** Seahorse measurement set up

| **Command** | **Time** |
|---|---|
| Baseline | Mix: 00:03:00<br>Wait: 00:00:00<br>Measure: 00:03:00<br>Cycle: 3 |
| Port A (compound) | Mix: 00:03:00<br>Wait: 00:00:00<br>Measure: 00:03:00<br>Cycle: 10 |
| Port B (oligomycin) | Mix: 00:03:00<br>Wait: 00:00:00<br>Measure: 00:03:00<br>Cycle: 3 |
| Port C (FCCP) | Mix: 00:03:00<br>Wait: 00:00:00<br>Measure: 00:03:00<br>Cycle: 3 |
| Port D (Rotenone/ Antimycin A) | Mix: 00:03:00<br>Wait: 00:00:00<br>Measure: 00:03:00<br>Cycle: 3 |

Supplementary Table 4**:** Primers used for Taqman RT-pPCR assay purchased from Thermo Fisher

| **Gene** | **Species** | **Assay ID** |
|---|---|---|
| LRP1 | Human | Hs01107912_g1 |
| LRP1B | Human | Hs01069153_m1 |

| | | |
|---|---|---|
| LRP2 | Human | Hs00218214_m1 |
| LRP4 | Human | Hs00391006_m1 |
| LRP8 | Human | Hs00182998_m1 |
| LDLR | Human | Hs01092524_m1 |
| TLR2 | Human | Hs02621280_s1 |
| UCP1 | Human | Hs01084772_m1 |
| CIDEA | Human | Hs00154455_m1 |
| FABP4 | Human | Hs01086177_m1 |
| TFAM | Human | Hs00273372_s1 |
| NRF1 | Human | Hs00602161_m1 |
| TBP | Human | Hs00427620_m1 |
| FIS1 | Human | Hs00211420_m1 |
| MFN1 | Human | Hs00966851_m1 |
| MFN2 | Human | Hs00208382_m1 |
| OPA1 | Human | Hs01047013_m1 |
| DNM1L | Human | Hs01552605_m1 |
| FASN | Human | Hs01005622_m1 |
| SREBF-1 | Human | Hs02561944_s1 |
| CD36 | Human | Hs00169627_m1 |
| GIPR | Human | Hs00609201_g1 |
| Serpina1a | Mouse | Mm02748447_g1 |
| Serpina1b | Mouse | Mm04207706_gH |
| Serpina1c | Mouse | Mm04207709_gH |
| Serpina1d | Mouse | Mm00842094_mH |

| | | |
|---|---|---|
| Serpina1e | Mouse | Mm00833655_m1 |
| Cidea | Mouse | Mm00432554_m1 |

Supplementary Table 5: Binding assays of Cerep panel Screening

| | **Assay** | **Source** | **Ligand** | **Conc** |
|---|---|---|---|---|
| **Receptors** | | | | |
| 1 | GABAB(B1b/B2) HumanGABA B GPCR Mass Spectrometry Binding (Antagonist Ligand) | Human recombinant (CHO cells) | CGP 54626 | 1 nM |
| 2 | $A_1$ (h) (antagonist radioligand) | Human recombinant (CHO cells) | $[^3H]$DPCP | 1 nM |
| 3 | $A_{2A}$ (h) (agonist radioligand) | Human recombinant (HEK-293 cells) | $[^3H]$CGS 21680 | 6 nM |
| 4 | $Alpha_{1A}$ (h) (antagonist radioligand) | Human recombinant (CHO cells) | $[^3H]$prazosin | 0.1 nM |
| 5 | $Alpha_{1B}$ (h) (antagonist radioligand) | Human recombinant (CHO cells) | $[^3H]$prazosin | 0.15 nM |
| 6 | $Alpha_{1D}$ (h) (antagonist radioligand) | Human recombinant (CHO cells) | $[^3H]$prazosin | 0.2 nM |
| 7 | $Alpha_{2A}$ (h) (antagonist radioligand) | Human recombinant (CHO cells) | $[^3H]$RX 821002 | 1 nM |
| 8 | $Alpha_{2B}$ (h) (antagonist radioligand) | Human recombinant (CHO cells) | $[^3H]$RX 821002 | 2.5 nM |
| 9 | $Beta_1$ (h) (agonist radioligand) | Human recombinant (HEK-293 cells) | $[^3H]$(-)CGP 12177 | 0.3 nM |
| 10 | $Beta_2$ (h) (agonist radioligand) | Human recombinant (CHO cells) | $[^3H]$(-)CGP 12177 | 0.3 nM |
| 11 | $AT_1$ (h) (antagonist radioligand) | Human recombinant (HEK-293 cells) | $[^{125}I][Sar^1,Ile^8]$-AT-II | 0.05 nM |

| | | | | |
|---|---|---|---|---|
| 12 | $B_2$ (h) (agonist radioligand) | Human recombinant (CHO cells) | [$^3$H]bradykinin | 0.3 nM |
| 13 | CB2 (h) (agonist radioligand) | Human recombinant (CHO cells) | [$^3$H]WIN 55212-2 | 0.8 nM |
| 14 | $CB_1$ (h) (agonist radioligand) | human recombinant (Chem-RBL cells) | [$^3$H]CP 55940 | 2 nM |
| 15 | $CCK_1$ ($CCK_A$) (h) (agonist radioligand) | Human recombinant (CHO cells) | [$^{125}$I]CCK-8s | 0.08 nM |
| 16 | $CCK_2$ ($CCK_B$) (h) (agonist radioligand) | Human recombinant (CHO cells) | [$^{125}$I]CCK-8s | 0.08 nM |
| 17 | $D_1$ (h) (antagonist radioligand) | Human recombinant (CHO cells) | [$^3$H]SCH 23390 | 0.3 nM |
| 18 | $D_{2S}$ (h) (agonist radioligand) | Human recombinant (HEK-293 cells) | [$^3$H]7-OH- DPAT | 1 nM |
| 19 | $D_{2L}$ (h) (antagonist radioligand) | Human recombinant (HEK-293 cells) | [$^3$H]methyl-spiperone | 0.3 nM |
| 20 | $ET_A$ (h) (agonist radioligand) | Human recombinant (CHO cells) | [$^{125}$I]endothelin-1 | 0.03 nM |
| 21 | $GABA_{A1}$ (h) (alpha1, beta2, gamma2) (agonist radioligand) | Human recombinant (CHO cells) | [$^3$H]muscimol | 15 nM |
| 22 | mGluR5 (h) (agonist radioligand) | Human recombinant (CHO cells) | [$^3$H]Quisqualate | 40 nM |
| 23 | CXCR2 (IL-8B) (h) (agonist radioligand) | Human recombinant (HEK-293 cells) | [$^{125}$I]IL-8 | 0.025 nM |
| 24 | CCR1 (h) (agonist radioligand) | Human recombinant (HEK-293 cells) | [$^{125}$I]MIP-1α | 0.01 nM |
| 25 | $H_1$ (h) (antagonist radioligand) | Human recombinant (HEK-293 cells) | [$^3$H]pyrilamine | 1 nM |
| 26 | H2 (h) (antagonist radioligand) | Human recombinant (CHO cells) | [$^{125}$I]APT | 0.075 nM |
| 27 | $CysLT_1$ ($LTD_4$) (h) (agonist radioligand) | Human recombinant (CHO cells) | [$^3$H]$LTD_4$ | 0.3 nM |

| | | | | |
|---|---|---|---|---|
| 28 | $MC_1$ (agonist radioligand) | Mouse endogenous (B16-F1 cells) | [$^{125}$I]NDP-α- MSH | 0.05 nM |
| 29 | $MC_4$ (h) (agonistradioligand) | Human recombinant (CHO cells) | [$^{125}$I]NDP-α- MSH | 0.05 nM |
| 30 | $M_1$ (h) (antagonist radioligand) | Human recombinant (CHO cells) | [$^3$H]pirenzepine | 2 nM |
| 31 | $M_2$ (h) (antagonist radioligand) | Human recombinant (CHO cells) | [$^3$H]AF-DX 384 | 2 nM |
| 32 | M3 (h) (antagonist radioligand) | Human recombinant (CHO cells) | [$^3$H]4-DAMP | 0.2 nM |
| 33 | $M_4$ (h) (antagonist radioligand) | Human recombinant (CHO cells) | [$^3$H]4-DAMP | 0.2 nM |
| 34 | $NK_1$ (h) (agonist radioligand) | Human endogenous (U373MG cells) | [$^{125}$I]-Substance PLYS3 | 0.05 nM |
| 35 | $Y_1$ (h) (agonist radioligand) | Human endogenous (SK-N-MC cells) | [$^{125}$I]peptide YY | 0.025 nM |
| 36 | N neuronal alpha4beta2 (h) (agonist radioligand) | Human recombinant (SH-SY5Ycells) | [$^3$H]cytisine | 0.6 nM |
| 37 | N muscle-type (h) (antagonist radioligand) | Human endogenous (TE671 cells) | [$^{125}$I]α-bungarotoxin | 0.5 nM |
| 38 | Delta (DOP) (h) (agonist radioligand) | Human recombinant (Chem-1 (RBL) cells) | [$^3$H]DADLE | 0.5 nM |
| 39 | kappa (h) (KOP) (agonist radioligand) | Human recombinant (RBL cells) | [$^3$H]U69593 | 0.5 nM |
| 40 | μ (MOP) (h) (agonist radioligand) | Human recombinant (HEK-293 cells) | [$^3$H]DAMGO | 0.5 nM |
| 41 | PPARgamma (h) (agonist radioligand) | Human recombinant (E. coli) | [$^3$H]rosiglitazone | 5 nM |
| 42 | PAF (h) (agonist radioligand) | Human recombinant (CHO cells) | [$^3$H]$C_{18}$-PAF | 1 nM |
| 43 | RARalpha (h) (agonist radioligand) | Human recombinant (insect cells) | 9-cis-Retinoic Acid [11,12-3H] | 1 nM |

| | | | | |
|---|---|---|---|---|
| 44 | 5-$HT_{1A}$ (h) (agonist radioligand) | Human recombinant (HEK-293 cells) | [$^3$H]8-OH-DPAT | 1 nM |
| 45 | 5-$HT_{1B}$ (h) (antagonist radioligand) | human recombinant (Chem-1 (RBL) cells) | [$^3$H]GR125743 | 1 nM |
| 46 | 5-$HT_{2A}$ (h) (agonist radioligand) | Human recombinant (HEK-293 cells) | [$^{125}$I](±)DOI | 0.1 nM |
| 47 | 5-$HT_{2B}$ (h) (agonist radioligand) | Human recombinant (CHO cells) | [$^{125}$I](±)DOI | 0.2 nM |
| 48 | 5-HT2C (h) (antagonist radioligand) | Human recombinant (HEK-293 cells) | [$^3$H]mesulergine | 1 nM |
| 49 | GR (h) (agonist radioligand) | Human endogenous (IM-9 cells) | [$^3$H]dexamethasone | 1.5 nM |
| 50 | ER Alpha Human Estrogen NHR Binding (Agonist radioligand) Assay, Cerep | Human recombinant | [$^3$H] Estradiol | 0.4 nM |
| 51 | PR (h) (agonist radioligand) | Human endogenous (T47D cells) | [$^3$H]progestero ne | 0.5 nM |
| 52 | AR(h) (agonist radioligand) | Human endogenous (LNCaP cells) | [$^3$H]methyltrien Olone | 1 nM |
| 53 | $V_{1a}$ (h) (agonist radioligand) | Human recombinant (CHO cells) | [$^3$H]AVP | 0.3 nM |
| **Ion channels** | | | | |
| 54 | Non-Selective Rat Glycine Ion Channel Strychnine Mass Spectrometry Binding Assay, Cerep | rat spinal cord | Strychnine | 10 nM |
| 55 | Glutamate (Non-Selective) Rat Ion Channel Glycine (Strychnine- Insensitive), Mass Spectrometry Binding Assay, Cerep | rat brain | MDL 105.519 | 2 nM |
| 56 | BZD (central) (agonist radioligand) | rat cerebral cortex | [$^3$H]flunitrazepam | 0.4 nM |

| | | | | |
|---|---|---|---|---|
| 57 | Cl- channel (GABA-gated) (TBOB site) (antagonistradioligand) | rat cerebral cortex | [$^3$H]TBOB | 3nM |
| 58 | AMPA (agonist radioligand) | rat cerebral cortex | [$^3$H]AMPA | 8 nM |
| 59 | Kainite (agonist radioligand) | rat cerebral cortex | [$^3$H]kainic acid | 5 nM |
| 60 | NMDA (antagonist radioligand) | rat cerebral cortex | [$^3$H]CGP39653 | 5 nM |
| 61 | PCP (antagonist radioligand) | rat cerebral cortex | [$^3$H]TCP | 10 nM |
| 62 | 5-HT3 (h) (antagonist radioligand) | Human recombinant (CHO cells) | [$^3$H]BRL 43694 | 0.5 nM |
| 63 | $Ca^{2+}$ channel (L, dihydropyridine site) (antagonist radioligand) | rat cerebral cortex | [$^3$H]nitrendipine | 0.25 nM |
| 64 | $Ca^{2+}$ channel (L, diltiazem site) (benzothiazepines) (antagonist radioligand) | rat cerebral cortex | [$^3$H]diltiazem | 15 nM |
| 65 | $Ca^{2+}$ channel (L, verapamil site) (phenylalkylamine) (antagonist radioligand) | rat cerebral cortex | [$^3$H]D888 | 3 nM |
| 66 | $Ca^{2+}$channel (N) (antagonist radioligand) | rat cerebral cortex | [$^{125}$I]ω-conotoxin GVIA | 0.001 nM |
| 67 | Potassium Channel hERG (human)- [3H] Dofetilide | Human recombinant (HEK-293 cells) | [$^3$H]Dofetilide | 3 nM |
| 68 | KV channel (antagonist radioligand) | rat cerebral cortex | [$^{125}$I]α-dendrotoxin | 0.01 nM |
| 69 | $Na^+$ channel (site 2) (antagonist radioligand) | rat cerebral cortex | [$^3$H]batrachotoxin | 10 nM |
| **Transporters** | | | | |
| 70 | Adenosine Guinea Pig Transporter Mass Spectrometry Binding (Antagonist Ligand) Assay, Cerep | Guinea-pig cerebral cortex | NBTI | 0.15 nM |

| | | | | |
|---|---|---|---|---|
| 71 | Norepinephrine transporter (h) (antagonist radioligand) | Human recombinant (CHO cells) | [$^3$H]nisoxetine | 1 nM |
| 72 | Dopamine transporter (h) (antagonist radioligand) | Human recombinant (CHO cells) | [$^3$H]BTCP | 4 nM |
| 73 | GABA transporter (antagonist radioligand) | rat cerebral cortex | [$^3$H]GABA (+ 10 µM isoguvacine) (+ 10 µM baclofen) | 10 nM |
| 74 | 5-HT transporter (h) (antagonist radioligand) | Human recombinant (CHO cells) | [$^3$H]imipramine | 2 nM |
| **Other enzymes** | | | | |
| 75 | MAO-A (antagonist radioligand) | rat cerebral cortex | [$^3$H]Ro 41-1049 | 10 nM |

Supplementary Table 6: Enzyme and uptake assays of Cerep panel screening

| | **Assay** | **Source** | **Substrates/ Stimulus/Tracer** |
|---|---|---|---|
| **Kinases** | | | |
| 1 | IRK (h) (InsR) | human recombinant | ATP + Ulight-Poly GAT [EAY (1:1:1)] (50 nM) |
| 2 | Lck kinase (h) | human recombinant (insect cells) | ATP + Ulight-Poly GAT [EAY (1:1:1)]n (25 nM) |
| 3 | PKCalpha (h) | human recombinant (insect cells) | ATP + CREBtide (CKRREI LSRRPSYRK) (20 nM) |
| **Other enzymes** | | | |
| 4 | COX1(h) | human recombinant | Arachidonic acid (3µM) + ADHP (25 µM) |
| 5 | COX2(h) | human recombinant (Sf9 cells) | Arachidonic acid (1.2 µM)+ ADHP (25 µM) |
| 6 | PDE3A *(h)* | human recombinant (Sf9 cells) | [$^3$H]cAMP + cAMP (0.5µM) |
| 7 | PDE4D2 (h) | human recombinant | [$^3$H]cAMP + cAMP (0.5µM) |

| | | | |
|---|---|---|---|
| | | (Sf9 cells) | |
| 8 | ACE (h) | human recombinant | Abz-FRK(Dnp)-P-OH (15 μM) |
| 9 | cathepsin G (h) | human neutrophils | N-succinyl-AAPF-pNa (1 mM) |
| 10 | acetylcholinesterase (h) | Human recombinant (HEK-293 cells) | Acetylthiocholine (400 μM) |
| 11 | MAO-B (h) recombinant enzyme | Human recombinant | D-Luciferin derivative (4 μM) |
| 12 | ATPase (Na+/K+) | Porcine cerebral cortex | ATP (2 mM) |

Supplementary Table 7**:** CRE-Luc2P reporter cell lines and their corresponding media

| **No #** | **Cell line** | **Receptor overexpression** | **Media** | **Supplement** |
|---|---|---|---|---|
| 1 | CHO CRE-luc2P | Human GIPR | Ham's F-12 Nutrient Mixture | 10% FBS, 50 μg/mL hygromycin B, and 250 μg/mL Zeocin |
| 2 | CHO CRE-luc2P | Mouse GIPR | | |
| 3 | HEK293 CRE-luc2P | No overexpression (empty) | DMEM (with high glucose/l-glutamine) | 10% FBS, 50 μg/mL hygromycin B |
